# IgG is differentially and selectively transferred across the placenta in HIV-infected women

**DOI:** 10.1101/479121

**Authors:** David R. Martinez, Youyi Fong, Shuk Hang Li, Fang Yang, Madeleine Jennewein, Joshua A. Weiner, Erin A. Harrell, Jesse F. Mangold, Ria Goswami, George Seage, Galit Alter, Margaret E. Ackerman, Xinxia Peng, Genevieve G. Fouda, Sallie R. Permar

**Author notes:** Lead authors: Corresponding author, (G.G.F) (S.R.P). Theses authors contributed equally (Genevieve G. Fouda, Sallie R. Permar).

## Abstract

**SUMMARY**

The transplacental transfer of maternal IgG to the developing fetus is critical for infant protection against infectious pathogens in the first year of life. However, factors that modulate the transplacental transfer efficiency of maternal IgG that could be harnessed for maternal vaccine design remain largely undefined. HIV-infected women have impaired placental IgG transfer, yet the mechanism underlying this impaired transfer is unknown, presenting an opportunity to explore factors that contribute to the efficiency of placental IgG transfer. We measured the transplacental transfer efficiency of maternal HIV and other pathogen-specific IgG in historical U.S. (n=120) and Malawian (n=47) cohorts of HIV-infected mothers and their HIV-exposed uninfected and HIV-infected infants. We then examined the role of maternal HIV disease progression, infant factors, placental Fc receptor expression, and IgG Fc region subclass and glycan signatures and their association with transplacental transfer efficiency of maternal antigen-specific IgG. We established 3 distinct phenotypes of placental IgG transfer efficiency in HIV-infected women, including: 1) efficient transfer of the majority of antigen-specific IgG populations; 2) generally poor IgG transfer phenotype that was strongly associated with maternal CD4+ T cell counts, hypergammaglobulinemia, and frequently yielded non-protective levels of vaccine-specific IgG; and 3) variable transfer of IgG across distinct antigen specificities. Interestingly, maternal IgG characteristics, such as binding to placentally expressed Fc receptors FcγRIIa and FcγRIIIa, IgG subclass frequency, and Fc region glycan profiles were associated with placental IgG transfer efficiency. These maternal IgG transplacental transfer determinants were distinct among different antigen-specific IgG populations. Our findings suggest that in HIV-infected women, both maternal disease progression and Fc region characteristics modulate the selective placental transfer of distinct IgG subpopulations, with implications for both the health of HIV-exposed uninfected infants and maternal vaccine design.

**Highlights:**

- Low peripheral blood CD4 + T cell count and hypergammaglobulinemia are associated with inefficient transplacental IgG transfer in HIV-infected women
- Antigen-specific IgG binding strength to placentally-expressed Fc receptors, but not placental Fc receptor expression levels, mediates selective placental IgG transfer
- Antigen-specific IgG Fc region glycan profiles also contribute to the selective placental IgG transfer of maternal IgG populations in HIV-infected women

## INTRODUCTION

Protective immunity in the first few months of life is reliant on maternal IgG that is passively transferred across the placenta (Dowling and Levy, 2014; Levy et al., 2013; Palmeira et al., 2012; Zash et al., 2016). This transplacental transfer of protective IgG can be enhanced by maternal vaccination during pregnancy. For example, it is estimated that worldwide incidence rates of neonatal tetanus decreased by 75% from the years 2000 to 2013 due to the wide-scale implementation of maternal tetanus toxoid vaccination during pregnancy (Khan et al., 2015). Furthermore, maternal immunization against influenza has also had a positive impact on infant health, with up to a 63% decrease in influenza infection rates in the first six months of life in infants born to vaccinated mothers compared to those born to unvaccinated women (Zaman et al., 2008). Yet, in 2015, despite the remarkable successes of maternal vaccination, >900,000 neonates died from vaccine-preventable respiratory infections worldwide (Liu et al., 2016). As an example, even though the maternal coverage for pertussis vaccination exceeds 80%, neonatal pertussis rates have increased three-fold over the last three decades in the U.S. alone (Healy et al., 2004). This may be partly due to the fact that maternal passively acquired pertussis-specific IgG can wane to low levels in newborns as early as two months of life, leaving the infant vulnerable to infection (Healy et al., 2004; Nunes et al., 2016). Therefore, there is an urgent need to 1) improve the transplacental IgG transfer efficiency of current maternal vaccines that are routinely administered during pregnancy and 2) develop novel maternal vaccine strategies designed for optimal placental IgG transfer to combat congenital and neonatal infections. On the other hand, minimizing the placental IgG transfer of monoclonal antibody therapies given to pregnant women for their own health is an important goal to improve the safety of this class of therapeutics during gestation.

The transplacental transfer of maternal IgG begins in the first trimester of pregnancy and by 20 weeks of gestation, maternal IgG levels in cord blood represent approximately 10% of the maternal blood levels (Malek et al., 1996). In normal pregnancies, by 37-40 weeks of gestation, infant cord blood levels can exceed maternal plasma IgG concentrations, often reaching levels >100% compared to those of their mothers (Kohler and Farr, 1966; Malek et al., 1996; Palmeira et al., 2012; Tatra and Placheta, 1979). Yet in the setting of specific maternal infections, such as HIV, the transplacental transfer of IgG is impaired (Dangor et al., 2015; de Moraes-Pinto et al., 1996; de Moraes-Pinto et al., 1993; de Moraes-Pinto et al., 1998a; de Moraes-Pinto et al., 1998b; Palmeira et al., 2012; Scott et al., 2005). Therefore, HIV-infected women represent a unique population to define factors that modulate the transplacental transfer of maternal IgG to the fetus. Moreover, HIV-exposed uninfected (HEU) infants have up to four-fold higher rates of morbidity and mortality from diarrheal and respiratory infections compared to unexposed infants (Dauby et al., 2016; Locks et al., 2017; Shapiro and Lockman, 2010; Shapiro et al., 2007; Slogrove et al., 2010; Weinberg et al., 2017). Several factors likely contribute to the high illness and death rates in HEU infants, including the poor transplacental transfer of protective maternal IgG (Adler et al., 2015; Brahmbhatt et al., 2006; Evans et al., 2016; Slogrove et al., 2016). Thus, understanding the mechanisms of impaired transplacental IgG transfer in HIV-infected women could inform strategies to improve the health of HEUs.

To reach the fetal circulatory system, maternal IgG must cross distinct placental cell barriers that make up the placental villous tree: the syncytiotrophoblast, the villous stroma, and fetal endothelial cells. The Fc receptor neonatal (FcRn) plays a key role in shuttling maternal IgG across the placenta to the fetal circulatory system (Roopenian and Akilesh, 2007; Simister, 2003; Simister and Mostov, 1989; Simister and Story, 1997). Yet, while syncytiotrophoblast cells express FcRn, neither stromal cells nor fetal endothelial cells express this canonical placental IgG shuttle receptor. Interestingly, other Fcγ receptors are also expressed in placental cells, yet their role in modulating the transplacental transfer of maternal protective IgG is unknown (Fouda et al., 2018; Kristoffersen and Matre, 1996; Martinez et al., 2018; Sedmak et al., 1991; Simister, 2003; Simister et al., 1996). Notably, Hofbauer cells located in the villous stroma express FcγRI and FcγRIII, and fetal endothelial cells – the last cell barrier crossed by maternal IgG before reaching the fetal circulatory system – express FcγRII (Kristoffersen et al., 1990; Martinez et al., 2018; Simister, 2003). Gaps in our understanding include how maternal IgG is transferred across this final placental cell barrier in the absence of FcRn, and whether FcγRI, FcγRII, or FcγRIII expression in placental cells contribute to the transplacental transfer of maternal IgG. In addition, IgG characteristics that impact Fc receptor (FcR) interactions, such as IgG subclass and/or Fc region glycans, could play a role in transplacental IgG transfer efficiency. The IgG subclass distribution among different antigen-specific IgG populations is distinct, and previous studies have indicated that this distribution impacts the transplacental transfer efficiency of different antigen-specific IgG populations (Ferrante et al., 1990).

In this study, we aimed to explore the mechanism(s) by which transplacental IgG transfer is impaired in HIV-infected women by identifying the determinants of placental IgG transfer through delineation of clinical and antibody characteristics that are associated with poor or efficient placental transfer of antigen-specific IgG populations in 167 HIV-infected pregnant women from the U.S and Malawi. We employed multivariable linear regression modeling to examine the association between transplacental IgG transfer and: 1) maternal HIV disease progression, 2) infant clinical factors, 3) placental factors such as FcR RNA expression, and 4) IgG Fc region characteristics including subclass, glycosylation profiles, and binding strength to placental FcRs. A deeper understanding of factors that modulate the transplacental transfer of maternal protective IgG will be important for improving infant health and extending the window of passively acquired IgG-mediated protection.

## RESULTS

### Clinical characteristics of U.S. and Malawian mother infant pairs

We assessed the transplacental transfer of HIV-specific and pathogen-specific IgG in two exiting cohorts of HIV infected women: the US-based Women and Infant transmission study (WITS) and the Malawian-based CHAVI009 study. The majority of the n=120 U.S. HIV infected women included in this study did not receive antiretroviral (ART) therapy during pregnancy but, ten of them (8.3%) received azidothymidine (AZT) monotherapy throughout pregnancy and two of them (1.6%) received AZT prophylaxis at delivery. In contrast, 100% of Malawian HIV-infected women received a single dose of nevirapine ART prophylaxis at delivery; and 13 of 47 (28%) of Malawian HIV-infected women with CD4+ T cell counts <250 (cells/mm^3^) received combination (Stavudine/Lamivudine/Nevirapine) antiretroviral (cART) therapy throughout pregnancy. Overall, HIV-infected U.S. women overall had lower plasma viral load and higher peripheral blood CD4+ T cell counts compared to HIV-infected Malawian women at delivery, despite more Malawian HIV-infected women being initiated on cART during pregnancy (Table 1). We measured total plasma IgG levels in U.S. and Malawian women as a surrogate measure of maternal hypergammaglobulinemia, a hallmark feature of advanced HIV-disease progression (Moir et al., 2001). HIV-infected U.S. women had lower total plasma IgG concentrations compared to Malawian women (median plasma IgG concentration levels of 16.3 mg/ml (range 2.5 – 123.5 mg/ml) vs 39 mg/ml (range 8.5 – 158.4 mg/ml) in U.S. vs Malawian women), with both populations having higher total plasma IgG concentrations compared to ~10mg/ml that is observed in normal pregnancies (Benster and Wood, 1970), consistent with HIV-associated hypergammaglobulinemia.

**Table 1.** Clinical characteristics of U.S. and Malawian HIV-infected women and their infants.

| Geographic location | Cohort | Mother infant pairs<br>N | Maternal ART treatment during pregnancy<br>N, (%) | Maternal ART treatment at delivery<br>N, (%) | Maternal peripheral blood CD4+ T cell count (cells/mm<sup>3</sup>) | Maternal plasma viral load (copies/ml) | Maternal total serum IgG (mg/ml) | HIV-exposed uninfected infants<br>N, (%) | HIV-infected infants<br>N, (%) | Infant gestational age (weeks) | Infant weight (Kg) | Infant sex |
| --- | --- | --- | --- | --- | --- | --- | --- | --- | --- | --- | --- | --- |
| U.S. | WITS | 120 | 10, (8.3%) | 2, (1.6%) | 642 (32-2,330) <sup>a</sup> | 33,929 (50-905,230) | 16.3 (2.5-123.5) | 120, (100%) | 0, (0%) | 40 (30-43) | 3.1 (1.4-4.5) | 41% F |
| Malawi | CHAVI009 | 47 | 13, (28%) | 47, (100%) | 381 (94-833) | 42,272 (400-343,721) | 39.9 (8.5-158.4) | 42, (89.3%) | 5, (10.6%) | 39 (33-44) | 3.1 (2.3-4.04) | 55% F |

<sup>a</sup> Median and range are shown

All 120 infants born to the U.S. HIV-infected women included in this study were uninfected, whereas five out of 47 (10.6%) infants born to Malawian HIV-infected women became infected with HIV *in utero* or peripartum. The U.S. HEU infants had a median gestational age of 40 weeks (range: 30-43 weeks), and Malawian HEU and infected infants had a median gestational age of 39 weeks (range: 33-44 weeks) (Table 1). There were no significant differences in infant birth weight, gestational age, or sex between the two populations.

### Distinct transplacental IgG transfer efficiency phenotypes in U.S. and Malawian HIV-infected women

Previous studies have demonstrated that the transplacental transfer efficiency of maternal vaccine antigen-specific IgG is impaired in the setting of maternal HIV infection (de Moraes-Pinto et al., 1996; de Moraes-Pinto et al., 1993). However, it is unclear if antigen-specific IgG populations are uniformly poorly transferred to the fetus in the setting of maternal HIV infection. We therefore measured HIV-specific IgG antibody levels in U.S. and Malawian paired mother-infant samples against various regions of the envelope glycoprotein including: gp120, variable-loop 3 (V3), variable-loop 1 and 2 (V1V2), and the gp41 membrane-proximal external region (MPER). We also measured antigen-specific IgG antibody levels against common neonatal pathogens, including: influenza hemagglutinin, pertussis toxin, tetanus toxoid, diphtheria toxin, rubella virus capsid, hepatitis B surface antigen, respiratory syncytial virus (RSV) F surface antigen, and *Haemophilus influenzae* type B (Hib) (Figure 1). Both HIV and standard neonatal pathogen-specific IgG responses were detectable at similar frequencies in both U.S. and Malawian HIV-infected women (Table S1).

**Figure 1.**
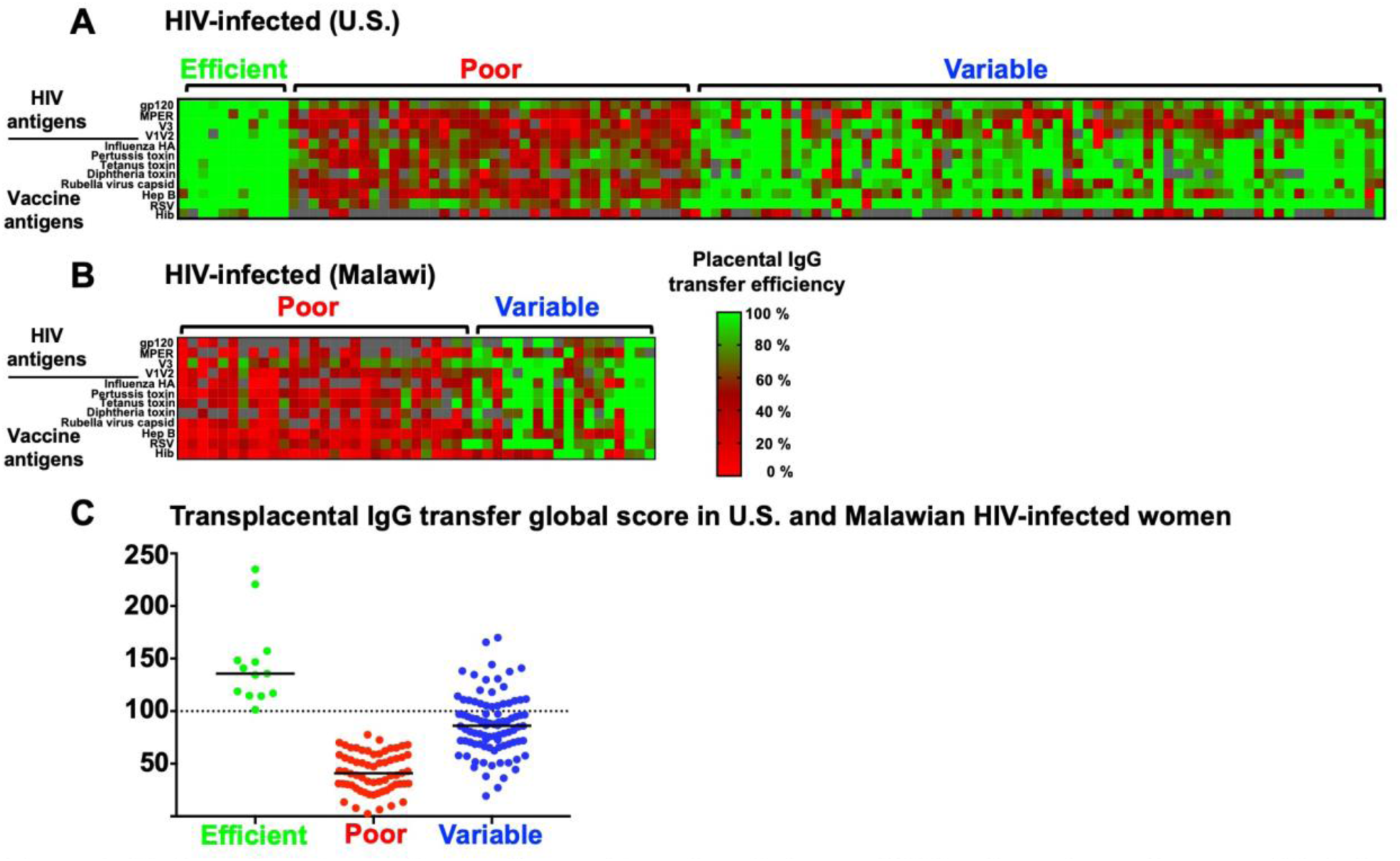
Distinct phenotypes of transplacental transfer efficiency of HIV and vaccine antigen-specific IgG among HIV-infected pregnant women. Transplacental IgG transfer efficiency in HIV-infected mother infant pairs from the (A) U.S. and (B) Malawi. (C) obalransplacental IgG transfer score among HIV-infected women defined to have efficient, poor, and variable transplacental IgG transfer. Bar denotes median. Grey squares denote uncalculated transfer ratios from concentrations outside the linear range.

In contrast to previous studies that have only reported a uniformly poor transplacental IgG transfer efficiency among antigen-specific IgG in HIV-infected women(de Moraes-Pinto et al., 1996; de Moraes-Pinto et al., 1998b), we observed three distinct transplacental transfer efficiency phenotypes, including: 1) efficient transplacental IgG transfer efficiency across the majority of measured antigen-specificities, 2) poor transplacental IgG transfer efficiency across the majority of antigen-specificities, and an unexpected phenotype of 3) variable transplacental IgG transfer efficiency across antigen-specific IgG populations (Figure 1).

To more quantitatively define transplacental IgG transfer efficiency phenotypes in HIV-infected women from both cohorts across antigen-specific IgG populations, we generated a global transplacental IgG transfer score for each mother-infant pair based on the mean transfer efficiency of the measured IgG populations (Figure 1). Using this scoring system, of 120 U.S., HIV-infected women, 8.3%, 30.8%, and 60.8% of mother-infant pairs were defined as exhibiting efficient, poor, and variable transplacental IgG transfer, respectively. In contrast, of 47 Malawian, HIV-infected women, 0%, 59.5%, and 40.5% mother-infant pairs were defined as exhibiting efficient, poor, and variable transplacental IgG transfer efficiency, respectively. We utilized the global transplacental IgG transfer score for U.S. and Malawian HIV-infected women using multivariable linear regression analyses to define maternal HIV disease progression factors that are associated with placental IgG transfer efficiency. We also generated separate transplacental IgG transfer efficiency scores for three specific IgG populations: gp120, tetanus toxoid, and pertussis toxin-specific IgG. We utilized the antigen-specific IgG transfer score of gp120, tetanus toxoid, and pertussis toxin-specific IgG in multivariable linear regression modeling to define IgG characteristics that are important for transplacental transfer efficiency of these specificities.

### U.S. and Malawian HIV-infected women with protective IgG concentrations passively transfer sub-protective levels of vaccine-specific IgG to their HEU infants

HEU infants are known to be more vulnerable to respiratory and diarrheal diseases and have higher rates of morbidity and mortality compared to HU infants (Afran et al., 2014; Filteau, 2009; Slogrove et al., 2016; Zash et al., 2016). We therefore examined the proportion of U.S. and Malawian infants that had cord blood plasma IgG levels below the protective threshold, despite being born to HIV-infected mothers with protective levels of vaccine antigen-specific IgG. We defined protective concentration thresholds as set by the World Health Organization (WHO) (Plotkin, 2010): tetanus toxoid-specific IgG (0.10 IU/ml), rubella-specific IgG (10 IU/ml), diphtheria toxin-specific IgG (0.10 IU/ml), and Hib-specific IgG (0.15 μg/ml). Of 120 U.S. HIV-infected women, 84%, 36%, 48%, and 22% had plasma concentrations above the protective level threshold for tetanus toxoid-specific IgG, rubella-specific IgG, diphtheria-specific IgG, and Hib-specific IgG, respectively (Figure 2). Yet, due to inefficient IgG transfer, 7%, 16%, 15%, and 41% of HEU infants born to these U.S. HIV-infected women with protective IgG levels displayed concentrations below the protective threshold for tetanus toxoid-specific IgG, rubella-specific IgG, diphtheria-specific IgG, and Hib-specific IgG, respectively. Similarly, in 47 Malawian HIV-infected women, 87%, 30%, 68%, and 21% had plasma concentrations above the protective level threshold for tetanus toxoid-specific IgG, rubella-specific IgG, diphtheria-specific IgG, and Hib-specific IgG, respectively. Yet, 15%, 71%, 50%, and 70% of infants born to these Malawian HIV-infected women with protective IgG levels had IgG concentrations below the protective threshold for tetanus toxoid, rubella, diphtheria, and Hib, respectively, highlighting the infant health consequences of impaired maternal IgG transfer in HIV-infected women on the infant health.

**Figure 2:**
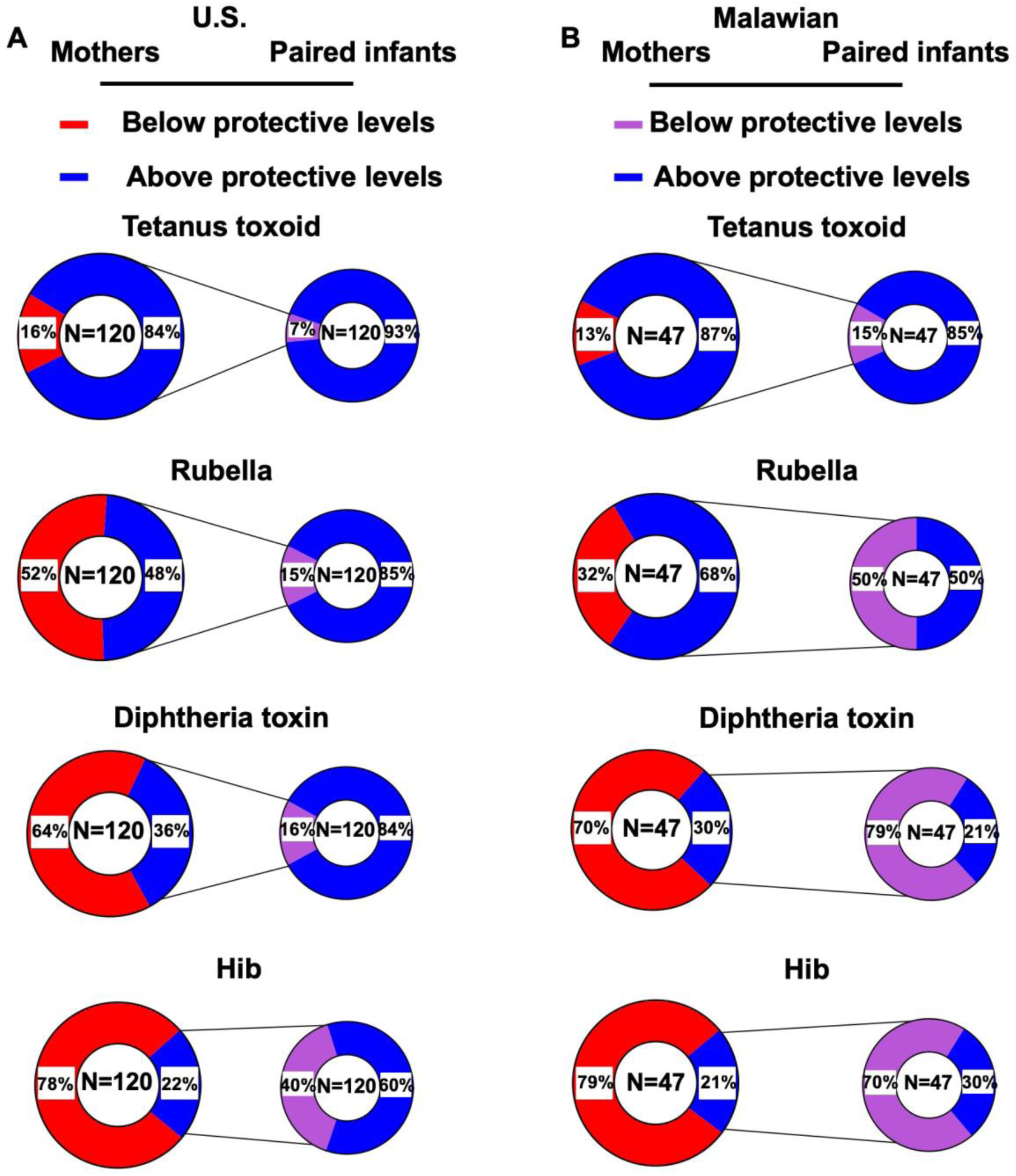
HEU infants receive suboptimal levels of protective maternal IgG against vaccine-preventable infections. Proportion of (A) U.S. and (B) Malawian mothers with levels of vaccine-specific IgG below (red) and above (blue) the established protective concentrations against tetanus toxoid, rubella, diphtheria toxin, and Haemophilus influenzae type B (Hib). Larger pie graphs show the proportion of mothers with IgG levels above and below the protective concentration. Smaller pie graphs show the proportion of infants born to women with protective IgG levels that fall below (purple) and above (blue) the protective IgG levels at birth.

### Maternal plasma antigen-specific IgG levels and transplacental IgG transfer efficiency in U.S. and Malawian women

To determine whether the antigen-specific IgG concentration in maternal plasma was a primary factor in the distinct transplacental IgG transfer efficiency of both HIV and vaccine antigen-specific IgG, we first examined the association between plasma antigen-specific IgG magnitude and transplacental IgG transfer efficiency (Table 2). In U.S. HIV-infected women, maternal plasma concentrations of HIV V3-specific and RSV surface F antigen-specific IgG was negatively associated with transplacental IgG transfer efficiency (slope: −0.24, p=0.05; slope: - 0.32, p<0.001, respectively). In Malawian HIV-infected women, tetanus toxoid-specific IgG was positively associated with transplacental transfer efficiency (slope: 0.50, p<0.004). Yet, maternal plasma antigen-specific IgG levels were not associated with transplacental transfer efficiency for the majority of HIV and vaccine antigen-specific IgG in both U.S. and Malawian HIV-infected women (Table 2), indicating that factors other than plasma antigen-specific IgG concentration contribute to the efficiency of transplacental IgG transfer.

**Table 2.** Associations between maternal plasma HIV and vaccine antigen-specific IgG levels and transplacental IgG transfer efficiency in HIV-infected pregnant women.

| Cohort | HIV<br>gp120 | HIV<br>MPER | HIV<br>V3 | HIV<br>V1V2 | Influenza<br>HA | Pertussis<br>toxin | Tetanus<br>toxin | Diphtheria<br>toxin | Rubella virus<br>capsid | Hep B | RSV | Hib |
| --- | --- | --- | --- | --- | --- | --- | --- | --- | --- | --- | --- | --- |
| U.S. | 0.08 (0.47) <sup>a</sup> | -0.07 (0.51) | **-0.24 (0.05)** | -0.06 (0.57) | -0.11 (0.26) | 0.08 (0.45) | -0.04 (0.67) | 0.16 (0.11) | 0.06 (0.57) | 0.02 (0.78) | **-0.32 (0.001)** | 0.21 (0.070) |
| Malawian | 0.00 (0.98) | -0.22 (0.29) | -0.13 (0.50) | -0.24 (0.11) | -0.07 (0.73) | 0.19 (0.18) | **0.50 (0.004)** | -0.10 (0.52) | -0.05 (0.74) | 0.05 (0.72) | 0.00 (0.99) | 0.12 (0.62) |

<sup>a</sup> Estimated regression slopes with p values in parentheses

### Maternal HIV disease progression factors and infant birth characteristics associations with transplacental IgG transfer efficiency

It remains unclear how maternal HIV disease progression factors contribute to the poor transplacental IgG transfer observed in HIV-infected women. Maternal CD4+ T cell counts were weakly positively correlated with transplacental IgG transfer efficiency in U.S. HIV-infected women (0.25 p<0.007) but not in Malawian HIV-infected women (0.10 p=0.47) (Figure S1). Interestingly, maternal total plasma IgG was negatively correlated with transplacental IgG transfer efficiency in both U.S. and Malawian HIV-infected women (Figure S1). We therefore sought to examine if maternal peripheral blood CD4+ T cell counts, plasma viral load, and total plasma IgG levels (or hypergammaglobulinemia) contribute to transplacental IgG transfer efficiency in both U.S. and Malawian women in a multivariable linear regression model. We examined maternal clinical characteristics and their relationship to the global transplacental IgG transfer score of each patient by multivariable linear regression. As maternal ART treatment during pregnancy has been associated with improved transplacental IgG transfer (Bosire et al., 2018), we first examined if maternal ART treatment during pregnancy in HIV-infected U.S., and Malawian HIV-infected women was associated with transplacental IgG transfer efficiency. Maternal ART monotherapy in U.S. HIV-infected women during pregnancy was not correlated with transplacental transfer efficiency (slope: 0.01, p=0.96, FDR=0.96) (Table 3). Similarly, combination ART treatment in Malawian HIV-infected women initiated during pregnancy was not associated with transplacental transfer efficiency (slope: 0.03, p=0.53, FDR=0.85). In this cohort, maternal peripheral blood CD4+ T cell counts were positively associated with transplacental IgG transfer efficiency in U.S. women (slope: 0.19, p<0.002, FDR<0.004), but not in Malawian women (slope: −0.03, p=0.85, FDR=0.85). In contrast, maternal plasma HIV RNA load was not associated with transplacental IgG transfer efficiency in either U.S. (slope: 0.02, p=0.74, FDR=0.93) or Malawian (slope: −0.05, p=0.80, FDR=0.85) women. Yet, maternal total plasma IgG concentrations were negatively associated with transplacental transfer efficiency in both U.S. (slope: −0.37, p=0.001, FDR=0.001) and in Malawian (slope: −0.40, p=0.005, FDR=0.02) HIV-infected women, indicating that hypergammaglobulinemia is associated with poor placental IgG transfer.

**Table 3.** Associations between maternal HIV disease progression and infant clinical characteristicsand transplacental IgG transfer efficiency.

| Maternal clinical factor/Infant characteristic at birth | U.S. |  |  | Malawian |  |  |
| --- | --- | --- | --- | --- | --- | --- |
|  | slope<sup>a</sup> | raw p | FDR<sup>b</sup> | slope | raw p | FDR |
| Maternal ART treatment during pregnancy | 0.01 | 0.96 | 0.96 | -0.03 | 0.53 | 0.85 |
| **Maternal peripheral blood CD4+ T cell count** | **0.19** | **0.002** | **0.004** | -0.03 | 0.85 | 0.85 |
| Maternal plasma HIV RNA load | 0.02 | 0.74 | 0.93 | -0.05 | 0.8 | 0.85 |
| **Maternal total plasma IgG level (hypergammaglobulinemia)** | **-0.37** | **0.001** | **0.001** | **-0.4** | **0.005** | **0.02** |
| Infant gestational age | 0.04 | 0.62 | 0.62 | 0.24 | 0.03 | 0.1 |
| Infant weight | 0.1 | 0.19 | 0.55 | -0.33 | 0.05 | 0.1 |
| Infant sex | -0.08 | 0.57 | 0.62 | 0.08 | 0.74 | 0.74 |

<sup>a</sup> Slope: Estimated regression slope values <sup>b</sup> FDR: false discovery rate adjusted p value Bolded values indicate statistical significance

Infant factors such as gestational age and birth weight have been associated with transplacental IgG transfer efficiency (Palmeira et al., 2012). To assess if these infant factors contributed to the distinct transplacental IgG transfer efficiency phenotypes, we examined infant gestational age, weight, and sex in a multivariable linear regression model in both U.S. HEU and Malawian HEU and HIV-infected infants (Table 3). In these primarily term infant cohorts, neither infant gestational age (U.S. slope: 0.04, p=0.62, FDR=0.62; Malawian slope: 0.24, p=0.03, FDR=0.1), nor weight (U.S. slope: 0.10, p=0.19, FDR=0.55; Malawian slope: −0.33, p=0.05, FDR=0.1), nor sex (U.S. slope: −0.08, p=0.57, FDR=0.62; Malawian slope: 0.08, p=0.74, FDR=0.74) were associated with transplacental transfer efficiency in either maternal-infant cohort.

### Maternal co-morbidities in Malawian HIV-infected women and transplacental IgG transfer efficiency

We next examined maternal co-morbidities that could potentially lead to altered placental IgG transfer such as maternal proteinuria (a marker of risk of preeclampsia) and syphilis co-infection, as defined by a maternal rapid plasma reagin (RPR) positive test in the Malawian cohort. These maternal co-morbidity data were not available for U.S. HIV-infected women. 8.5% of Malawian HIV-infected women tested positive for the RPR test, whereas 36% tested positive for trace levels of proteinuria. Interestingly, proteinuria, in addition to total plasma IgG concentration, was associated with efficient transplacental IgG transfer in Malawian HIV-infected women (slope: 0.55, p=0.01). In contrast, maternal co-infection with syphilis was not significantly associated with the transplacental transfer efficiency of maternal IgG (slope: −0.79, p=0.06).

### Placental FcR expression and transplacental IgG transfer

To examine the role of placental FcR expression on the transplacental transfer efficiency of maternal IgG, we performed RNAseq analysis on available placental biopsy tissues only from Malawian HIV-infected women (n=44), as placental biopsy tissues were not collected in the U.S. cohort. Maternal placental FcRn, FcγRIIa, FcγIIb, FcγIIc, FcγIIIa, and FcγIIIb mRNA was detectable in all Malawian HIV-infected women (Figure S2), yet the levels of expression as defined by Log2 copies per million (CPM) were variable (Figure S3). We then compared the levels of placental FcR mRNA expression in HIV-infected women with variable and poor transplacental transfer of maternal IgG. Placental FcRn, FcγRIIa, FcγIIb, FcγIIc, and FcγIIIa expression levels strongly correlated with each other, whereas FcγIIIb had lower overall expression levels and a weak correlation with FcRn, FcγRIIa, FcγIIb, FcγIIc, and FcγIIIa expression levels (Figure S2). We did not observe statistically significant differences in FcR expression levels in HIV-infected women with variable and poor transplacental IgG transfer efficiency (Figure S3). We examined Fc receptor expression levels and their relationship to the global transplacental IgG transfer score of each patient by multivariable linear regression. There was no significant association between placental FcR expression levels and transplacental IgG transfer efficiency.

### Binding of maternal antigen-specific IgG for placental FcRs and transplacental IgG transfer

We next assessed the binding of maternal plasma antigen-specific antibodies to placentally-expressed FcRs and their common polymorphic variants in Malawian HIV-infected women (n=47). We measured maternal gp120, tetanus toxoid, and pertussis toxin-specific IgG binding to FcRn, FcγRIIb, FcγRIIIb, the high affinity polymorphic forms FcγRIIa 131H and FcγRIIIa 158V, and the low affinity polymorphic variants FcγRIIa 131R and FcγRIIIa 158F, as well as to complement protein C1q (Figure S4). The binding magnitude of gp120 and tetanus toxoid-specific IgG strongly correlated among the measured placental FcRs, whereas the binding magnitude of pertussis toxin-specific IgG to placental FcRs exhibited variable correlations among FcRs (Figure S5). After correcting for predictors of transplacental IgG transfer in HIV-infected Malawian women (total plasma IgG levels and proteinuria), we examined if gp120, tetanus toxin, and pertussis toxin-specific IgG binding to placentally expressed FcRs was associated with transplacental IgG transfer efficiency. We related Fc receptor binding magnitude of gp120, tetanus toxin, and pertussis toxin-specific IgG to their unique antigen-specific transplacental IgG transfer score by multivariable linear regression. Neither maternal gp120 nor pertussis toxin-specific IgG binding to placentally expressed FcRs was associated with their transplacental IgG transfer efficiency (Table 4). In contrast, maternal tetanus toxoid-specific IgG binding to all FcRs positively, albeit weakly, associated with transplacental IgG transfer efficiency (slope: 0.23, p<0.05) (Table 4). Moreover, binding of tetanus toxoid-specific IgG to FcγRIIa 131H (slope: 0.18, p<0.03), FcγRIIa 131R (slope: 0.15, p<0.03), and FcγRIIIa 158F (0.21, p<0.05) were positively associated with transplacental IgG transfer efficiency of this specificity (Table 4), suggesting distinct FcR interaction determinants of placental IgG transfer efficiency for different antigen-specific IgG populations.

**Table 4.**
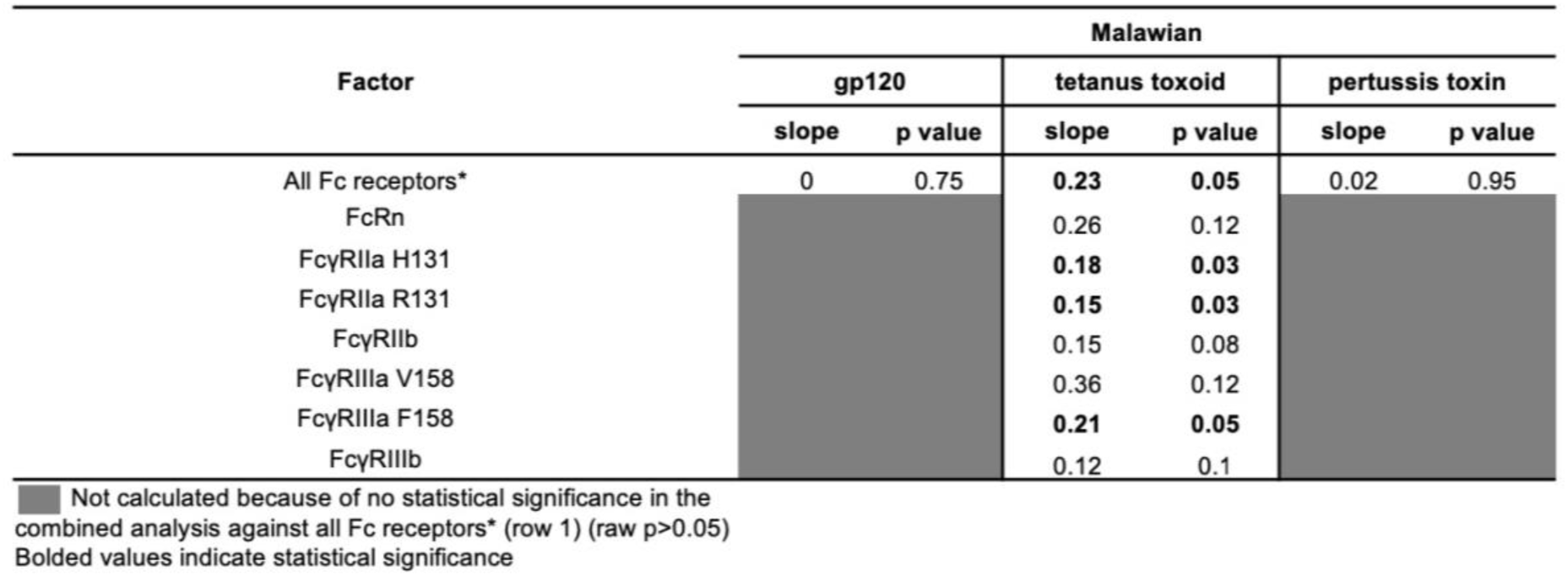
Associations of transplacental IgG transfer efficiency and maternal plasma antigen-specific IgG binding to acentally-expressed Fc receptors in Malawian women.

### IgG subclass and transplacental IgG transfer efficiency in HIV-infected women with variable transplacental IgG transfer efficiency

To further explore IgG Fc characteristics that play a role in antigen-specific IgG transplacental transfer efficiency, we focused on three antigen-specific IgG populations of HIV-infected women defined to have variable transplacental IgG transfer efficiency phenotype: gp120, tetanus toxoid, and pertussis toxin-specific IgG. We measured gp120, tetanus toxoid, and pertussis toxin-specific IgG subclass distribution in both U.S. (n=50) and Malawian (n=16) HIV-infected women with variable transfer of antigen-specific IgG populations (Figure S6). We focused on defining IgG characteristics of gp120-specific IgG because it is generally poorly transferred and on tetanus toxoid and pertussis toxin-specific IgG because they are generally efficiently transferred (Figure S6). The frequency of gp120, tetanus toxoid, and pertussis toxin-specific IgG1 subclass responses was higher than that of the other IgG subclasses in both U.S. and Malawian HIV-infected women. Gp120, tetanus toxoid, and pertussis toxin-specific IgG2 subclass responses were less frequently detected in all antigen-specific populations in U.S. compared to Malawian women (U.S.: gp120-specific 3%, tetanus toxoid-specific 2%, and pertussis toxin-specific 3%; Malawian: gp120-specific 6%, tetanus toxoid-specific 25%, and pertussis toxin-specific 6%, Figure S6). Similarly, U.S. HIV-infected women had relatively lower frequencies of gp120, tetanus toxoid, and pertussis toxin-specific IgG3 subclass responses compared to Malawian women (U.S.: gp120-specific 17%, tetanus toxoid-specific 10%, and pertussis toxin-specific 7%; Malawian: gp120-specific 25%, tetanus toxoid-specific 44%, and pertussis toxin-specific 13%, Figure S6). Lastly, U.S. HIV-infected women also had overall lower frequencies of detectable gp120, tetanus toxoid, and pertussis toxin-specific IgG4 subclass responses compared to Malawian HIV-infected women (U.S.: gp120-specific 2%, tetanus toxoid-specific 27%, and pertussis toxin-specific 2%; Malawian: gp120-specific 19%, tetanus toxoid-specific 81%, and pertussis toxin-specific 0%, Figure S6). In contrast to U.S. HIV-infected women, Malawian HIV-infected women were boosted with a tetanus vaccine during the third trimester of pregnancy, perhaps explaining the overall higher frequencies of tetanus toxoid-specific IgG subclass responses.

We then related IgG subclass frequency of gp120, tetanus toxin, and pertussis toxin-specific IgG to their unique antigen-specific transplacental IgG transfer score using multivariable linear regression modelling. We assessed the isolated contribution of antigen-specific IgG subclass distribution to placental IgG transfer efficiency using a multivariable linear regression model that corrected for identified predictors (maternal CD4+ T cell counts and total plasma IgG concentrations) of transplacental IgG transfer. Due to the high collinearity of IgG1 subclass and antigen-specific IgG response frequency, the frequency of antigen-specific IgG responses was included instead of IgG1 subclass in the model (Table 5). Interestingly, in U.S. women, the frequency of maternal gp120-specific IgG3 subclass responses (slope: −0.68, p<0.03), the frequency of maternal tetanus toxoid-specific IgG4 subclass responses (slope: −0.52, p=0.04), and the frequency of maternal pertussis-specific IgG responses (slope: −0.33, p=0.02) were negatively associated with their transplacental IgG transfer (Table 5). We did not find that IgG subclass distribution of gp120, tetanus toxoid, or pertussis toxin-specific IgG, to be associated with their transplacental IgG transfer efficiency in Malawian women, likely due to the smaller cohort size.

**Table 5.** Associations of transplacental IgG transfer efficiency and antigen-specific IgG subclass responses in U.S and Malawian HIV-infected women with variable transplacental IgG transfer.

| Cohort | Maternal IgG subclass | U.S. |  |  |  |  |  |
| --- | --- | --- | --- | --- | --- | --- | --- |
|  |  | gp120-specific IgG transfer |  | tetanus toxoid-specific IgG transfer |  | pertussis toxin-specific IgG transfer |  |
|  |  | slope | p value | slope | p value | slope | p value |
| U.S. | Antigen-specific IgG frequency <sup>b</sup> | 0.03 | 0.82 | -0.09 | 0.41 | **-0.33** | **0.02** |
|  | IgG2 frequency | 1.78 | 0.08 | 0.78 | 0.15 | 0.70 | 0.44 |
|  | IgG3 frequency | **-0.68** | **0.03** | 0.09 | 0.81 | -0.27 | 0.61 |
|  | IgG4 frequency | LF <sup>a</sup> | LF | **-0.52** | **0.04** | 0.15 | 0.87 |
| Malawian | Antigen-specific IgG frequency | -0.79 | 0.28 | 0.35 | 0.11 | LF | LF |
|  | IgG2 frequency | -0.45 | 0.63 | 0.09 | 0.84 | 0.83 | 0.39 |
|  | IgG3 frequency | 0.28 | 0.74 | 0.18 | 0.64 | 0.52 | 0.22 |
|  | IgG4 frequency | 0.12 | 0.87 | -0.02 | 0.97 | -0.25 | 0.80 |

<sup>a</sup> LF: low frequency. <sup>b</sup> Due to the high collinearity of IgG1 subclass responses and antigen-specific IgG response magnitude, the frequency of IgG1 subclass was not included in the combined model of IgG characteristics Bolded values indicate statistical significance

### IgG Fc region glycans and transplacental IgG transfer efficiency in HIV-infected women with variable transfer efficiency

To examine the potential contribution IgG Fc region glycan profiles in mediating transplacental IgG transfer efficiency, we measured the composition of Fc region glycans in the overall poorly transferred gp120-specific IgG population, and generally efficiently transferred tetanus toxoid and pertussis toxin-specific IgG populations in U.S. HIV-infected women with variable transplacental IgG transfer (n=50) (Figure 3A). We did not observe statistically significant differences of agalactosylated (G0) (Figure 3B) Fc region glycans of gp120, tetanus toxoid, and pertussis toxin-specific IgG. In contrast, we observed statistically significantly different profiles of monogalactosylated (G1) (p<0.0001, Figure 3C), digalactosylated (G2) (p<0.01, Figure 3D), fucosylated (p<0.0001, Figure 3E), bisected (p<0.003, Figure 3F), disialylated (p<0.0001, Figure 3G), mono-sialylated (p<0.0001, Figure 3H), and total sialylated (p<0.0001, Figure 3I) Fc region glycans between the isolated IgG for the same specificities. To determine if specific Fc region glycans were associated with transplacental IgG transfer efficiency, we performed a principal component analysis in which we examined the proportional variance of each principal component as it related to the transplacental IgG transfer efficiency. Principal component 1 (composed of G0, G1, G2, mono sialic acid, and total sialic acid, Figure 4, Table S2) accounted for 59-64% of the variance of transplacental IgG transfer of gp120, tetanus toxoid, and pertussis toxin-specific IgG (Table S2), whereas principal component 2 (composed of fucose, disialic acid, and bisecting, Figure 4, Table S2) accounted for 16-24% of the variance of transplacental IgG transfer for the same specificities.

**Figure 3:**
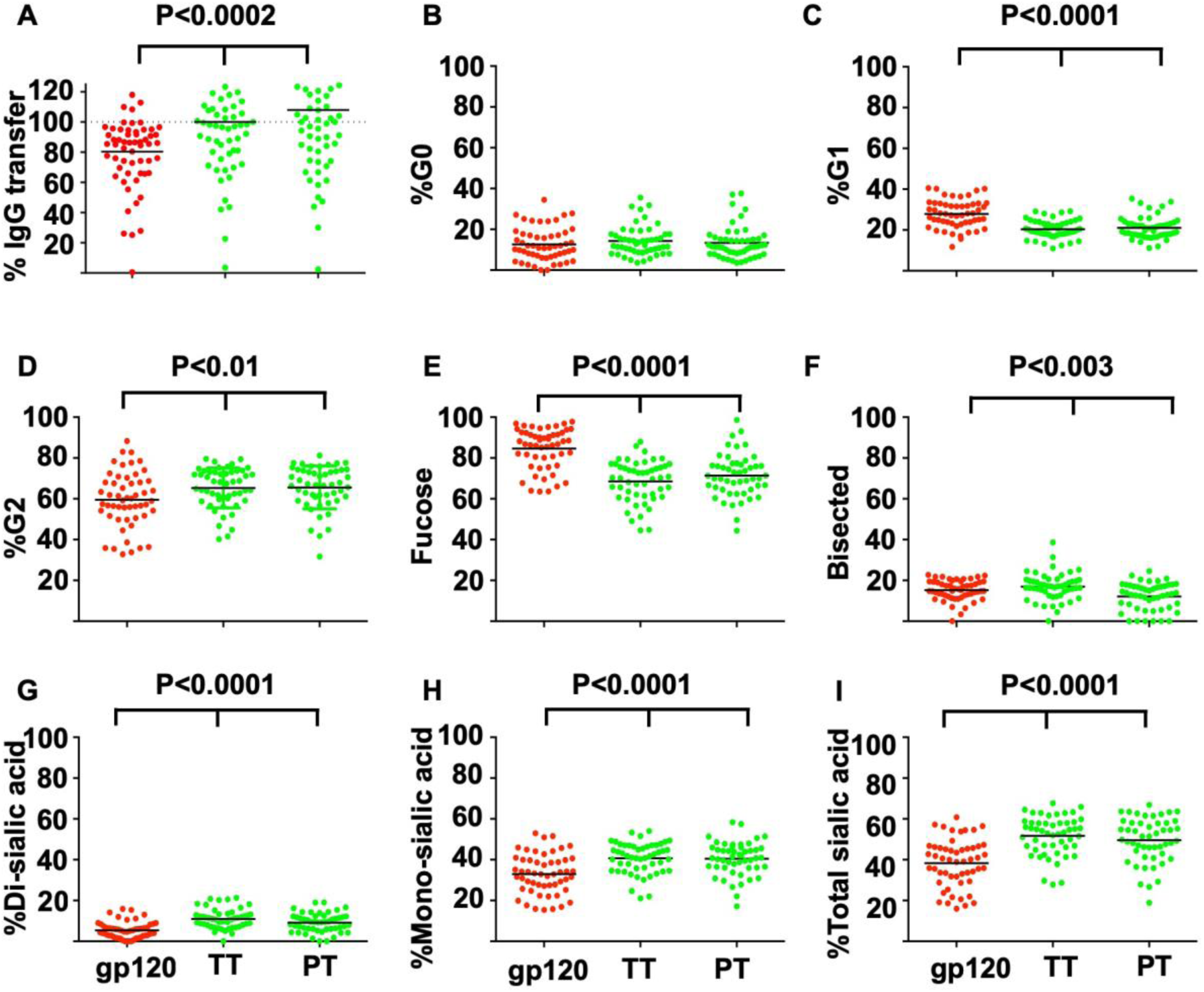
Differences in Fc region glycan profiles of gp120, tetanus toxoid, andpertussis toxin-specific IgG in U.S. HIV-infected women with variable transplacental IgG transfer. (A) . transplacental transfer efficiency of gp120,tetanus toxoid (TT), and pertussis toxoid (PT)-specific IgG. Frequency of (B) GO, (C) G1, (D) G2, (E) fucose, (F) bisected, (G) di-sialic acid, (H) mono-sialic acid, (I) total sialic acid of Fc region glycans of gp120, tetanus toxoid, and pertussis toxin-specific IgG. Displayed is p value from a Kruskal Wallis test, dotted line denotes 100.transfer.

**Figure 4:**
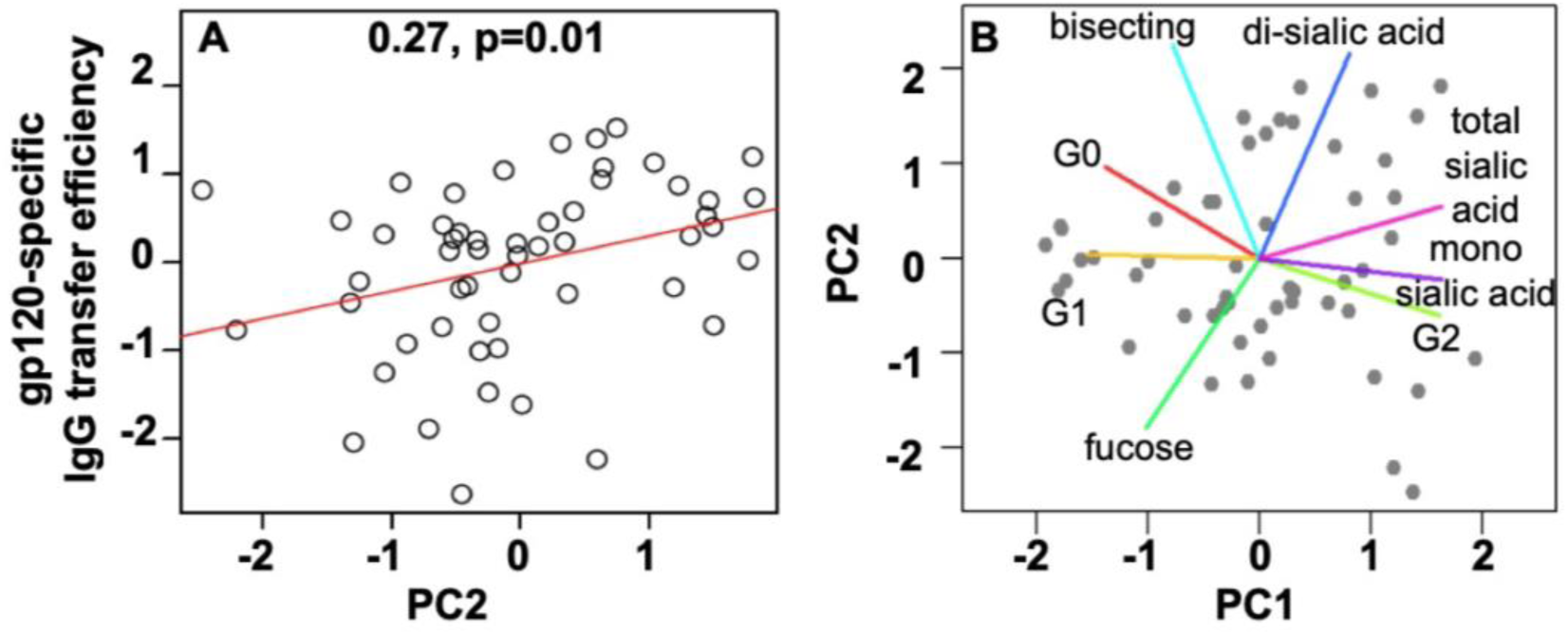
gp120-specific IgG Fc glycan profile are associatedwith transplacental IgG transfer efficiency in U.S. HIV-infected women with variable transplacental IgG transfer. Regressionmodeling of gp120-specific IgG glycan (A) PC2 (Fucose, Bisecting,and Di-sialic acid) and transplacental IgG transfer efficiency. (B) Fcglycan PC 1 and 2 loadings plot analysis of individual Fc region glycan profiles of gp120-specific IgG.

Previous studies reported that a lower frequency of Fc region fucose glycans led to increased binding strength to FcγRIIIa, suggesting that Fc region glycans may modulate binding to Fc receptors (Masuda et al., 2007; Niwa et al., 2004; Okazaki et al., 2004). Interestingly, poorly transferred gp120-specific IgG had higher frequencies of Fc region fucose, whereas efficiency transferred tetanus toxoid and pertussis toxin had overall higher levels of fucose (Figure 3D), suggesting that Fc region glycans may modulate the selective transfer of IgG subpopulations. We therefore related Fc region glycan profiles of gp120, tetanus toxin, and pertussis toxin-specific IgG to their unique antigen-specific transplacental IgG transfer score by multivariable linear regression. We fit a multivariable linear regression model that corrected for identified clinical predictors of transplacental IgG transfer efficiency (maternal CD4+ T cell counts and total plasma IgG concentrations). We found that Fc region glycan signatures of tetanus toxoid or pertussis toxin-specific IgG were not associated with their transplacental IgG transfer efficiency. In contrast, gp120-specific IgG Fc region fucose, bisecting, and disialic acid glycan frequencies (PC2) were weakly positively associated with transplacental IgG transfer efficiency of this specificity (slope: 0.27, p<0.01) (Figure 4A, Figure 4B, and Table 6), suggesting that Fc region fucose, bisected, and disialylated glycans can modulate transplacental IgG transfer efficiency of some but not all antigen-specific IgG subpopulations. We next examined the isolated contribution of each individual gp120-specific IgG Fc region glycan as it related to transplacental IgG transfer efficiency. We observed weak yet significant associations with individual gp120-specific Fc region glycans (Table 6). Gp120-specific IgG Fc region fucose and di-sialic acid frequencies were significantly but very weakly associated with transplacental IgG transfer efficiency, suggesting a collective effect of these Fc region glycans on placental IgG transfer (Table 6).

**Table 6.**
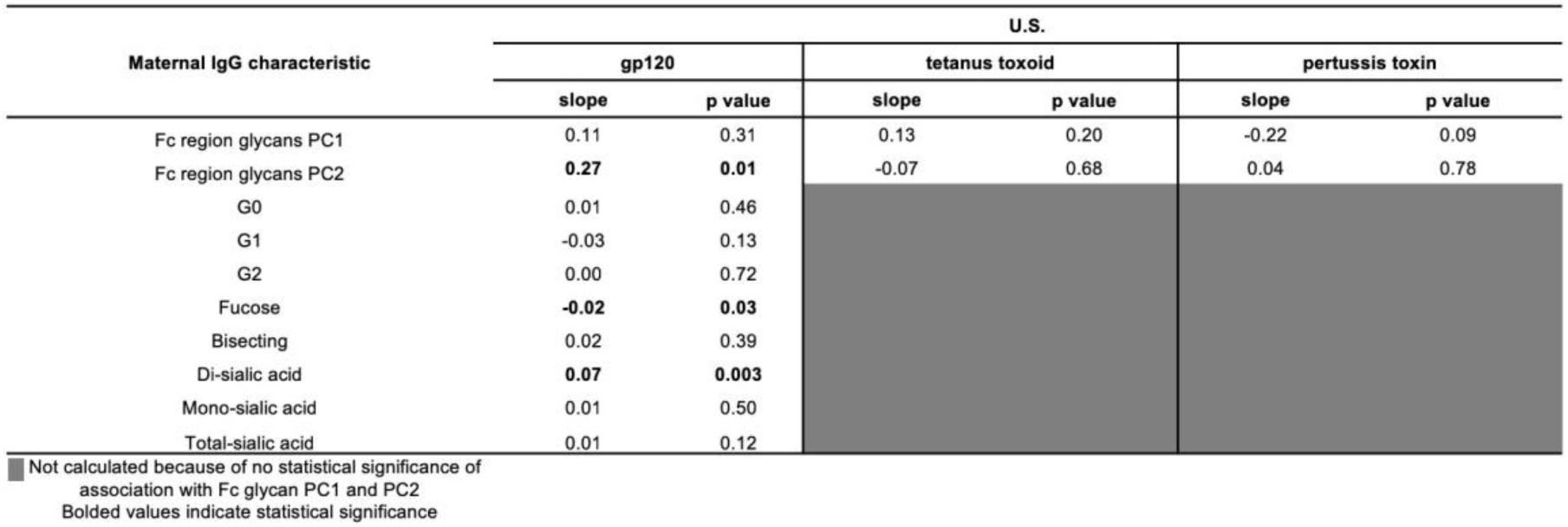
Associations transplacental IgG transfer efficiency and antigen-specific plasma IgG levels and Fc region glycan profiles in U.S. women with variable transplacental IgG transfer.

| Maternal IgG characteristic | U.S. |  |  |  |  |  |
| --- | --- | --- | --- | --- | --- | --- |
|  | gp120 |  | tetanus toxoid |  | pertussis toxin |  |
|  | slope | p value | slope | p value | slope | p value |
| Fc region glycans PC1 | 0.11 | 0.31 | 0.13 | 0.20 | -0.22 | 0.09 |
| Fc region glycans PC2 | **0.27** | **0.01** | -0.07 | 0.68 | 0.04 | 0.78 |
| G0 | 0.01 | 0.46 |  |  |  |  |
| G1 | -0.03 | 0.13 |  |  |  |  |
| G2 | 0.00 | 0.72 |  |  |  |  |
| Fucose | **-0.02** | **0.03** |  |  |  |  |
| Bisecting | 0.02 | 0.39 |  |  |  |  |
| Di-sialic acid | **0.07** | **0.003** |  |  |  |  |
| Mono-sialic acid | 0.01 | 0.50 |  |  |  |  |
| Total-sialic acid | 0.01 | 0.12 |  |  |  |  |

■ Not calculated because of no statistical significance of association with Fc glycan PC1 and PC2 Bolded values indicate statistical significance

### Antigen-specific IgG magnitude and transplacental IgG transfer efficiency in HIV-infected women with variable transfer efficiency

After again correcting for identified maternal predictors of transplacental IgG transfer efficiency, we reexamined the isolated role of antigen-specific plasma IgG concentrations and their association with transplacental IgG transfer efficiency in U.S. women with variable transplacental IgG transfer efficiency. We examined antigen-specific magnitude of gp120, tetanus toxin, and pertussis toxin-specific IgG by multivariable linear regression using their unique antigen-specific transplacental IgG transfer score. Maternal gp120-specific IgG concentrations were significantly associated with transplacental transfer efficiency in HIV-infected women with variable transplacental IgG transfer efficiency (slope: 0.38, p=0.01) (Table 6). In contrast, neither maternal tetanus toxoid-specific IgG concentrations (slope: −0.04, p=0.78) nor maternal pertussis toxin-specific IgG concentrations (slope: −0.03, p=0.89) were associated with their transplacental IgG transfer efficiency in U.S. HIV-infected women with variable transplacental IgG transfer.

### Combined analysis of IgG characteristics associated with transplacental IgG transfer efficiency in HIV-infected women with variable transfer efficiency

Finally, to examine the combined impact of IgG characteristics on transplacental transfer efficiency, we fit a linear regression model that combined each of the identified characteristics of gp120, tetanus toxoid, and pertussis toxin-specific IgG found to be associated with placental IgG transfer efficiency in U.S. HIV-infected women with variable transplacental IgG transfer efficiency. We related these IgG characteristics to their unique antigen-specific transplacental IgG transfer score by multivariable linear regression. Again, due to the high collinearity of IgG1 subclass responses and antigen-specific IgG response magnitude, the frequency of IgG1 subclass was not included in the combined model of IgG characteristics (Table 7). The frequency of tetanus toxoid-specific IgG subclass IgG4 responses was also negatively associated with transplacental IgG transfer of tetanus toxoid-specific IgG (slope: −0.65, p<0.01) (Table 7). The high frequency of IgG4 subclass responses and the negative association with transplacental IgG transfer was only observed for tetanus toxoid-specific IgG. Moreover, in the combined model of IgG Fc characteristics, the frequency of IgG2 subclass responses was strongly positively associated with transplacental IgG transfer efficiency of gp120, tetanus toxoid, and pertussis toxin-specific IgG (slope: 1.21, p<0.01, slope: 0.99, p<0.01, slope: 1.26, p<0.01, respectively). In addition, magnitude of maternal plasma gp120-specific IgG was positively associated with transplacental IgG transfer efficiency this specificity (slope: 0.27, p<0.03). Finally, gp120- specific IgG Fc region glycan profiles that comprised PC2 remained weakly positively associated with transplacental IgG transfer (slope: 0.26, p<0.01). Altogether, these findings suggest that IgG Fc characteristics differentially mediate the selective placental transfer efficiency of distinct IgG subpopulations.

**Table 7.**
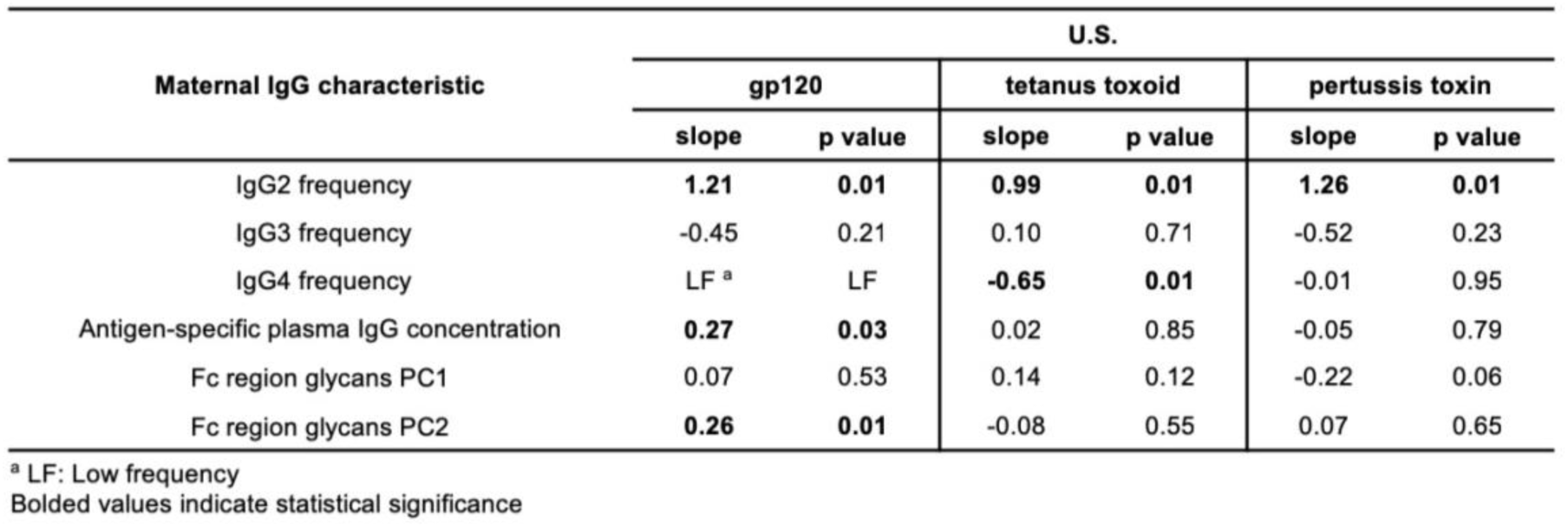
Combined analysis of maternal IgG characteristics associated with transplacental IgG transfer efficiency in U.S HIV-infected women with variable transplacental IgG transfer efficiency.

## DISCUSSION

A major roadblock for improving the protection of newborns via maternal passively-acquired maternal IgG is our limited understanding of the molecular interactions of IgG and placental FcRs that modulate transplacental IgG transfer. Placental IgG transfer is impaired in the setting of certain maternal infections, such as HIV and malaria (Cumberland et al., 2007; de Moraes-Pinto et al., 1996; de Moraes-Pinto et al., 1993; de Moraes-Pinto et al., 1998b; Palmeira et al., 2012). Thus, we sought to define factors that associated with poor or efficient placental IgG transfer in the pathologic setting of maternal HIV-infection. Interestingly, we observed wide variability in the efficiency of transplacental transfer of antigen-specific IgG in U.S. and Malawian HIV-infected women who received minimal ART treatment (Figure 1). In U.S. HIV-infected mother infant pairs, we observed efficient, poor, and variable transplacental IgG transfer. In contrast, in Malawian HIV-infected mother infant pairs, we only observed poor and variable transplacental IgG transfer. It was surprising to observe the variable transplacental IgG transfer efficiency phenotype in the majority of HIV-infected women, and it suggests that the placental transfer is distinctly regulated for different antigen-specific IgG populations. Malawian HIV-infected women had more evidence of advanced HIV disease, with higher plasma viral load, lower peripheral blood CD4 + T cell counts, and higher concentrations of total plasma IgG compared to HIV-infected women from the U.S. (Table 1), potentially accounting for the lack of an efficient transplacental IgG transfer efficiency in this cohort.

Previous studies have reported the suboptimal transplacental transfer of protective IgG in HIV-infected women (Abu-Raya et al., 2016; Gupta et al., 2014; Jones et al., 2011), raising the possibility that the poor transplacental transfer of maternal protective IgG may contribute to the higher rates of infectious diseases in HEU infants. Accordingly, we observed that between 7- 79% of infants born to HIV-infected women with protective plasma IgG titers against tetanus, rubella, diphtheria, and Hib, were born with IgG concentrations below protective thresholds against these common neonatal pathogens (Figure 2). Interestingly, despite having a tetanus toxoid booster vaccine during pregnancy, Malawian HIV-infected women still had overall poor transplacental IgG transfer of tetanus toxoid-specific IgG. Moreover, up to 15% of HEU infants born to Malawian HIV-infected women were below the protective titer for tetanus-specific IgG at birth, suggesting that maternal vaccination alone may not be sufficient to overcome the poor transplacental IgG transfer in this population. These findings highlight that the poor transplacental transfer of maternal protective IgG in HIV-infected pregnant women puts their HEU infants at risk for vaccine-preventable infectious diseases.

In our study, biomarkers of maternal HIV disease progression, including CD4+ T cell counts and hypergammaglobulinemia, were associated with poor placental IgG transfer. However, maternal plasma HIV load was not associated with transplacental IgG transfer efficiency in either U.S. or Malawian HIV-infected women. This finding contrasts with a prior study that reported maternal plasma HIV load and poor transplacental IgG transfer efficiency in a cohort of 50 HIV, clade A virus-infected Kenyan women (Farquhar et al., 2005). This disparate finding could be due to differences in maternal HIV disease progression and potentially HIV clade-specific differences. Total IgG concentrations in maternal plasma above a certain high threshold (greater than ~15mg/ml) have previously been associated with poor transplacental IgG transfer efficiency (Gonçalves et al., 1999; Hartter et al., 2000; Michaux et al., 1966; Okoko et al., 2001; Palmeira et al., 2012), and accordingly, maternal hypergammaglobulinemia was associated with poor transplacental IgG transfer efficiency in both U.S. and Malawian HIV- infected women (Table 3). In addition, Malawian HIV-infected women had higher median concentrations of plasma total IgG concentrations compared to U.S. HIV-infected women (Table 1). Altogether, our findings suggest that low maternal peripheral blood CD4+ T cell counts and hypergammaglobulinemia negatively impact the transplacental transfer of maternal IgG. While it is possible that high maternal plasma IgG concentrations are a biomarker of advanced HIV-disease progression that tracks together with low peripheral blood CD4 + T cell counts in these primarily untreated cohorts, it is also possible that maternal high plasma IgG concentrations lead to impaired transplacental IgG transfer by saturating placentally expressed FcRs that shuttle maternal IgG to the fetus (Englund, 2007; Palmeira et al., 2012; Wilcox et al., 2017). In addition, it is possible that the B cell dysfunction that leads to hypergammaglobulinemia in HIV-infected women also results in altered Fc region characteristics (i.e., altered IgG subclass composition or Fc region glycan profiles) that lead to impaired transplacental transfer of some IgG subpopulations.

While the role of placentally expressed FcRn in mediating the transplacental transfer of maternal IgG is clear (Palmeira et al., 2012; Roopenian and Akilesh, 2007; Simister and Mostov, 1989), the role of noncanonical placental FcRs in transplacental transfer of maternal IgG remains unexplored (Simister, 2003; Simister and Story, 1997). We did not detect overall placental FcR expression differences in HIV-infected, Malawian women with distinct phenotypes of transplacental IgG transfer (Figure S2, Figure S3), suggesting the determinants of IgG transfer efficiency are more likely to be modulated by IgG characteristics or other factors, even though it remains unclear if cell-type-specific FcR abundances may play a role. Moreover, differences in the overall placental FcRs expression levels in Malawian HIV-infected women were not associated with transplacental IgG transfer efficiency. Interestingly, the binding of tetanus toxoid-specific IgG to placentally expressed FcγRIIa 131H, FcγRIIa 131R, and FcγRIIIa F158, positively associated with transplacental IgG transfer efficiency in Malawian HIV-infected women, suggesting that IgG interaction with noncanonical placental FcRs may play a role in mediating the transplacental transfer of some antigen-specific IgG populations (Table 4). The distinct outcomes in FcR binding and transplacental transfer efficiency among maternal gp120, tetanus toxoid, and pertussis toxin-specific IgG, may be explained by differences in IgG characteristics among these distinct antibody populations. For example, tetanus toxoid-specific IgG responses in Malawian HIV-infected women had higher frequencies of IgG4 subclass responses compared to both gp120 and pertussis-specific IgG responses (Figure S6). Furthermore, gp120, tetanus toxoid, and pertussis-specific IgG exhibited distinct Fc region glycan profiles (Figure 3) which could contribute to the lack of a uniform binding strength to distinct placentally-expressed Fc receptors (Figure S2).

Maternal IgG subclass is known to be an important factor in transplacental IgG transfer (Fouda et al., 2018; Palmeira et al., 2012), with IgG1 and IgG4 being the most efficiently transferred (Garty et al., 1994). In this cohort of U.S. HIV-infected women, the frequency of gp120-specific IgG3 subclass, tetanus toxoid-specific IgG1, and pertussis-specific IgG4 subclass responses negatively associated with transplacental transfer efficiency (Table 5), suggesting that IgG subclass frequency of some antigen-specific IgG populations may negatively impact the transplacental IgG transfer. Moreover, maternal IgG2 subclass frequency of gp120, tetanus toxoid, and pertussis toxin-specific IgG strongly positively associated with efficient transplacental IgG transfer in the combined model (Table 7) of IgG characteristics (Table 5).

However, the association of gp120, tetanus toxoid, and pertussis toxin-specific IgG subclass prevalence and efficient transplacental IgG transfer should be interpreted with caution given the low frequency of IgG2 subclass responses in U.S. HIV-infected women with variable transplacental IgG transfer efficiency. We did not find the same association of IgG subclass frequencies and transplacental IgG transfer in Malawian HIV-infected women, potentially attributable to a smaller cohort size or more advanced HIV disease progression. It is also possible that higher frequency of IgG1 and IgG4 subclass responses track together with hypergammaglobulinemia in HIV-infected women. As it has been reported that IgG subclasses have distinct affinity for FcRn, FcγRII, and FcγRIII (Abdiche et al., 2015; Bruhns et al., 2009), IgG subclass-specific affinity to placental FcRs could be an important, yet unexplored, determinant of transplacental IgG transfer efficiency.

The Fc region glycosylation profile at the conserved glycosylation site may also be an important factor that impacts transplacental IgG transfer efficiency, due to its role in mediating IgG binding to certain placental Fc receptors including: FcγRI, FcγRII, and FcγRIII. While the affinity of IgG Fc to the canonical IgG placental shuttle receptor, FcRn, is not likely dependent on this Fc glycosylation profile (Martin et al., 2001), previous studies have reported differences in IgG Fc region glycans in mother infant pairs (Williams et al., 1995), suggesting that IgG Fc glycan-dependent placentally-expressed Fcγ receptors may modulate transplacental IgG transfer efficiency. While one study that examined IgG Fc region glycans in ten mother-infant pairs found no significant differences in Fc region glycosylation patterns in maternal and infant IgG populations (Einarsdottir et al., 2013), a more recent study found overall IgG glycan differences in maternal IgG and those that were passively transferred to the infant (Jansen et al., 2016).

However, these studies examined glycosylation profiles in total maternal IgG as opposed to that of antigen-specific IgG populations, potentially masking distinctions in placentally transferred Fc region glycan profiles. Consistent with a previous study that reported distinct Fc region glycans profiles of antigen-specific IgG populations (Mahan et al., 2016), poorly transplacentally-transferred gp120-specific IgG had overall differences in the relative amounts of certain Fc region glycans compared to that of efficiently-transferred tetanus and pertussis toxoid-specific IgG (Figure 4). Further, we identified that the Fc region glycan profile was associated with transplacental IgG transfer efficiency of gp120-specific IgG (Figure 4, Table 6). Fucosylated, bisected, and disialylated IgG Fc region glycans were weakly associated with efficient transplacental IgG transfer efficiency of gp120-specific IgG in combination, but had limited contribution individually (Table 6), suggesting a cooperative effect among Fc region glycans. Fc region fucose glycans have been shown to mediate binding strength to FcγRIIIa in vitro (Okazaki et al., 2004). Accordingly, we also found that poorly transferred gp120-specific IgG had lower frequencies of Fc region fucose, suggesting that Fc region glycans modulate transplacental IgG transfer efficiency, likely through altered binding to placental Fc receptors such as FcγRIIIa. Our findings suggest that antigen-specific IgG Fc profiles modulate the selective placental transfer efficiency of distinct IgG subpopulations.

While we aimed to comprehensively examine factors that mediate the transplacental IgG transfer in HIV-infected women, our study has some limitations. Both cohorts lacked detailed vaccination history, with the exception of tetanus vaccination during pregnancy in Malawian women. Another limitation of our study is the lack of a robust clinical record for U.S. HIV-infected women of potential co-morbidities that could also impact the efficiency of transplacental IgG transfer. Nonetheless, we examined the role of clinical signs of preeclampsia and maternal syphilis in Malawian HIV-infected women and designed our linear regression models to correct for these factors that are known to affect transplacental IgG transfer. Moreover, while we found that placental FcR RNA expression levels were not distinct among HIV-infected Malawian women with variable and poor transplacental IgG transfer, it is known that gene copy numbers may vary within the FcR locus and could complicate the interpretation of FcR mRNA expression levels (Hollox and Hoh, 2014; Lassauniere and Tiemessen, 2016). Our inability to detect FcR RNA expression level differences could also be due to the absence of an efficient transplacental IgG transfer group in HIV-infected Malawian women. Another caveat to our study is that factors in U.S. HIV-infected women that were associated with transplacental IgG transfer efficiency were not always similarly correlated with transplacental IgG transfer efficiency in Malawian HIV-infected women. While these differences could be explained by differences in cohort sizes, health, and/or genetic differences in U.S. and Malawian HIV-infected women, our findings should be validated in larger cohorts of HIV-infected mother infant pairs. It should also be noted that the calculated transplacental IgG transfer score was specifically designed to detect differences among HIV-infected U.S. and Malawian women with distinct IgG transfer phenotypes and was inherently arbitrary. Finally, as only 5 of 167 infants were HIV infected, our study was not designed to identify immune correlates of protection against mother to child transmission of HIV, but instead was geared towards defining determinants of transplacental IgG transfer efficiency.

Altogether, our findings reveal that the placental transfer of maternal IgG is a selective process, particularly in the setting of HIV infection. Our results suggest that a combination of factors, including IgG FcR binding strength, subclass, and glycan profiles including fucose, but not placental FcR expression levels or antigen-specific IgG concentration, collectively play a role in the selective and differential placental transfer of maternal antigen-specific IgG (Figure 5). Our results also suggest that the transplacental transfer efficiency is differentially regulated by distinct Fc region characteristics across antigen-specific IgG subpopulations. These findings also shed light on the potential design of strategies to improve the transplacental IgG transfer to the fetus via Fc region modifications of some antigen-specific IgG (i.e., tetanus toxoid and pertussis toxin-specific IgG). Moreover, our findings have important implications for improving the transplacental IgG transfer achieved by routinely administered vaccine-elicited IgG responses in pregnancy such as the tetanus, diphtheria, and pertussis vaccine (i.e., Tdap), especially in HIV-infected women. For example, future studies could explore adjuvant-mediated modulation of the Fc region characteristics of vaccine-elicited IgG to increase binding to placentally-expressed FcRs or drive specific Fc region glycosylation profiles with the goal of improving transplacental IgG transfer efficiency. These data also suggest that maternal HIV treatment that prevents CD4+ T cell loss and reduces maternal hypergammaglobulinemia is likely to improve the transplacental transfer of maternal protective IgG, which should be studied in combination ART-treated HIV-infected maternal populations. Future work that further defines the determinants of placental IgG transport will ultimately inform strategies to improve the transfer of maternal IgG to the vulnerable fetus, extending the window of maternal IgG-mediated infant protection in the first year of life.

**Figure 5:**
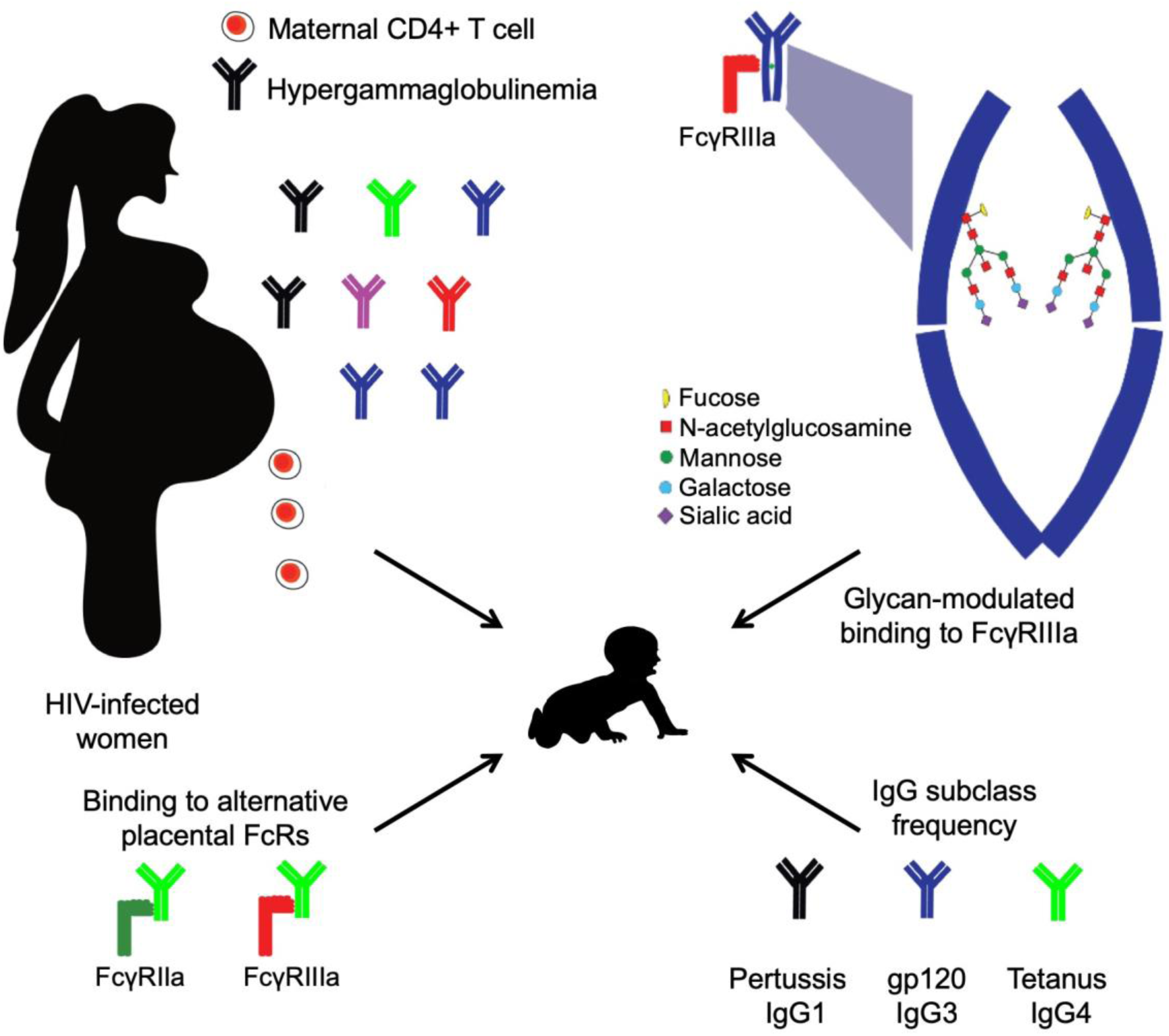
Model of the selective placental transfer of maternal IgG in HIV-infected pregnant women.

## EXPERIMENTAL PROCEDURES

### Study population

HIV-infected women and their HIV-infected or HIV-exposed uninfected (HEU) infants were selected from two previously described maternal-infant cohorts based on the availability of maternal plasma and paired infant cord blood plasma sample from delivery (Fouda et al., 2013; Permar et al., 2015; Sacha et al., 2015). N=120 maternal and infant cord-blood samples from delivery were obtained from the Woman and Infant Transmission Study (WITS) cohort. WITS study participants were HIV-infected women from North America, thus primarily infected with clade B virus. All 120 infants included from the WITS cohort were HEU as determined by plasma HIV load PCR testing at delivery. Forty-seven maternal and infant cord blood plasma samples from delivery were obtained from the Center for HIV/AIDS Vaccine Immunology 009 (CHAVI009) cohort of HIV-infected women from Malawi. Mothers and infants enrolled in the CHAVI009 study received a single dose Nevirapine at delivery. Five infants from the CHAVI009 cohort were HIV-infected *in utero,* and n=42 infants were HEU as determined by whole blood HIV DNA PCR testing at delivery.

### Ethics statement

Approval was obtained from the institutional review board at each collaborating institution and enrollment clinical sites to utilize de-identified maternal and infant cord blood plasma samples from the WITS and CHAVI009 cohorts.

### Measurement of antigen-specific IgG levels

A binding antibody multiplex assay (BAMA) was used to measure maternal and infant HIV and non-HIV-specific IgG responses. Briefly, carboxylated beads were coupled to HIV and non-HIV antigens as described previously (Tomaras et al., 2008). A total of 5 × 10^6^ carboxylated fluorescent beads (Luminex Corp) were covalently coupled to 25µg of proteins and peptides tested. Recombinant HIV and non-HIV antigens included: Con6gp120, MNnegD11gp120, gp70MNV3, gp70BcaseAV1V2, MulVgp70_his, Bio-V3.C (Bio-KKKNNTRKSIRIGPGQTFYATGDIIGDIRQAHC), Bio-MPER03 (Bio-KKKNEQELLELDKWASLWNWFDITNWLWYIR), Bio-MPER656, (Bio-KKKNEQELLELDKWASLWNWFNITNWLW), Tetanus toxoid (Pfenex, Inc), Pertussis toxin (Sigma-Aldrich), Influenza hemagglutinin/A/Solomon Islands/03/2006 (Protein Sciences Corp), Rubella virus capsid (Abcam), Diphtheria toxin (Sigma), *Hemaphilus Influenza B* (Hib)Type B oligosaccharide-Human Serum Albumin conjugate (BEI Resources), Hepatitis B surface antigen (Abcam), and Respiratory syncytial virus (RSV) F surface antigen (DSCAV-1) (a kind gift of Dr. Barney Graham, NIH VRC). Peptides were purchased from CPC Scientific (Sunnyvale, CA). Maternal and infant plasma was tested against Con6gp120, MNnegD11gp120, gp70MNV3, gp70BcaseAV1V2, MulVgp70_his, Bio-V3.C, Bio-MPER03, Bio-MPER656, Tetanus toxoid, Pertussis toxin, Influenza hemagglutinin/A/Solomon Islands/03/2006, Rubella virus capsid, Diphtheria toxin, and RSV DSCAV-1 surface F antigen at 1:100 dilution in serum diluent (1X PBS, 1% milk, 5% normal goat serum, 0.05% Tween-20). Maternal and infant plasma was tested against Hib Type B oligosaccharide-Human Serum Albumin conjugate and Hepatitis B surface antigen at a 1:25 dilution in a modified serum diluent to reduce background (1X PBS, 1% milk, 5% normal goat serum, 0.05% Tween-20, 0.05% polyvinyl alcohol, and 0.08% polyvinylpyrrolidone). Maternal and infant plasma HIV and vaccine antigen-specific IgG was detected with a mouse anti-human IgG (Southern Biotech, Birmingham, AL) phycoerythrin-conjugated antibody at 2µg/ml as described previously (Tomaras et al., 2008). Hyperimmuneglobulin isolated from HIV-seropositive donors (HIVIG) standard was used to calculate concentration of Con6gp120-specific IgG antibodies in mother infant pairs. A membrane proximal external region (MPER)-specific IgG (2F5) was used to calculate the concentration of MPER-specific IgG in mother infant pairs. A V3-specific IgG (CH22) was used to calculate the concentration of maternal and infant V3-specific IgG. A V1V2-specific IgG (CH58) was used to calculate the concentration of V1V2-specific IgG in mother infant pairs. An influenza hemagglutinin-specific IgG (CH65) was used to calculate the concentration of flu-specific IgG in mother infant pairs. Finally, an RSV surface antigen-specific mAb (Palivizumab - MedImmune) was used to calculate the concentration of RSV-specific IgG in mother infant pairs. Concentration of tetanus toxoid, pertussis toxin, rubella virus capsid, *Haemophilus influenzae* type B, hepatitis B, and diphtheria toxin-specific IgG was determined by using commercially available WHO international standards (National Institute for Biological Standards and Controls – anti-tetanus toxoid sera: 26/288, anti-pertussis sera: 06/140, anti-rubella sera: 91/688, anti-Haemophilus influezae type B: 09/222, anti-Hepatitis B: 07/164, anti-diphtheria toxin: 10/262). We measured protective concentration thresholds in U.S. and Malawian HIV-infected women as set by the World Health Organization (WHO) (Plotkin, 2010): tetanus toxoid-specific IgG (0.10 IU/ml), rubella-specific IgG (10 IU/ml), diphtheria toxin-specific IgG (0.10 IU/ml), and Hib-specific IgG (0.15 μg/ml). Normal human sera samples from uninfected donors were used as a negative control for HIV antigens. All MFI values were blank bead and well subtracted. All mother-infant samples were tested at the same dilution. Antibody measurements were acquired using a Bio-Plex 200 instrument (Bio-Rad).

### Calculation of transplacental IgG transfer efficiency

Maternal IgG transplacental transfer efficiency was calculated as infant antigen-specific IgG concentration/maternal antigen-specific IgG concentration X 100. Efficient transplacental transfer was defined as >99% transfer efficiency for the majority (>75%) of measured antigen-specific IgG. Variable transplacental transfer was defined as efficient (>99%) and poor (<60%) transfer efficiency of measured antigen-specific IgG within a mother infant pair. Poor transplacental transfer was defined as <60% transfer efficiency for the majority (>75%) of measured antigen-specific IgG.

### Measurement of HIV and non-HIV antigen-specific IgG subclass

To measure HIV and vaccine-antigen IgG subclass responses, maternal samples were tested against a Con6gp120, MNgDneg11gp120, gp70MNV3, gp70BcaseAV1V2, tetanus toxoid, pertussis toxin, MulVgp70_his, influenza hemagglutinin/A/Solomon Islands/03/2006 (Protein Sciences Corp), and a blank bead at a 1:40 dilution by BAMA. Biotinylated antibody reagents specific for each IgG subclass were used at 4 µg/ml: anti-IgG1-biotin (BD Pharmingen, clone: G17-1), anti-IgG2-biotin (BD Pharmingen, G18-21), anti-IgG3-biotin (Calbiochem, clone: HP6047), and anti-IgG4-biotin (BD Pharmingen, clone: G17-4). Plates were washed 3X and streptavidin-conjugated phycoerythrin antibody diluted was used at 1:100 dilution and added to wells. All MFI values were blank subtracted.

### IgG Fc region glycosylation analysis

Maternal plasma samples were heat inactivated at 56°C for 1 hour. Maternal plasma samples were incubated with 25µl of uncoated streptavidin coated magnetic beads (New England Biolabs) at room temperature (RT) in rotation. MNnegD11gp120, tetanus toxoid, and pertussis toxin antigens were biotinylated with LC-LC-biotin (Thermo Scientific) according to the manufacturer’s protocol. Streptavidin coated magnetic beads were activated with 0.5M NaCl, 20mM Tris-HCl [pH 7.5], 1mM EDTA and incubated with biotinylated antigens at RT for 1 hour in rotation. Maternal plasma and coupled beads were mixed and incubated at RT for 1 hour in rotation. Adsorbed plasma samples were removed and antigen-specific IgG bound to the antigen-coupled beads was digested with IDEZ (New England Biolabs) at 37°C for 1 hour in rotation. Cleaved Fc region supernatants were digested with PNGaseF (New England Biolabs) according to manufacturer specifications. To precipitate Fc region glycans, ice-cold ethanol was added and incubated at −20°C for 30 min. Plates were spun and the ethanol layer was transferred to clean plates. Plates were spun in a Centrivap (Labconco Corporation) and dried glycans were labeled with APTS dye (Life Technology) in Sodium Cyanoborohydride in Tetrahydrofuran (Sigma). Labeled glycans were transferred to filtered plates (Harvard Apparatus) containing P-2 slurry (Bio-Rad) and excess APTS dye was removed by washing with water. Cleaned Fc region labeled glycans were run immediately on a 3130 Genetic analyzer Sanger sequencer (Applied Biosystems) as described previously (Mahan et al., 2015).

### Measurement of total plasma IgG

Goat anti-human polyclonal IgG (Life Technologies) was used to coat high binding 384 well ELISA plates (Corning) with 3µg/ml overnight at 4°C. Plates were washed and blocked with SuperBlock (4% whey protein, 15% goat serum, and 0.5% Tween 20 diluted in 1X PBS) for 2 hours at RT. Maternal plasma IgG was tested at 1:1000 starting dilution and was serially diluted in 3-fold dilutions. Reagent grade normal human serum IgG (Sigma) was used as a standard at a starting concentration of 10µg/ml and was serially diluted in 3-fold dilutions. An anti-human IgG HRP conjugated antibody produced in goat (Sigma Aldrich) was used at 1:10,000 dilution. After washing the plates, SureBlue reserve TMB substrate (KPL) was added. Substrate reactions were stopped by adding an equal volume of Stop solution (KLP). Optical densities were read at 450nm using a Spectramax plate reader. OD values within the linear range of a 5-parameter standard curve were used to interpolate the total IgG concentration of plasma samples.

### Placental RNAseq analysis and assessment of FcR expression levels

Placental biopsy samples in RNAlater from HIV-infected Malawian women were thawed on ice. Fifty mg of tissue was homogenized in Trizol (Invitrogen), and total RNA was purified with miRNeasy extraction kit (Qiagen) according to the manufacturer guidelines. RNA quality was assessed on Bioanalyzer 2100 instrument using RNA 6000 Nano Kit (Agilent). Total RNA libraries were prepared from 500 ng purified RNA using TruSeq Stranded Total RNA with Ribo-Zero Gold Kit (Illumina). Deep sequencing was performed on a Nextseq500 sequencer (Illumina) using 75bp paired-end reads. Raw BCL (base call) files generated from NextSeq sequencer were converted to FASTQ files using bcl2fastq Conversion Software v2.18. During BCL to FASTQ processing, bcl2fastq also separates multiplexed samples and removes adapters. Each pair of a FASTQ file was sequentially mapped to human ribosomal RNA and hemoglobin sequences using a gapped aligner STAR (Dobin et al., 2013) to make sure that rRNA was depleted and there was no hemoglobin contamination. Raw RNA-seq data in FASTQ file format was quality controlled during and after sequencing to identify potential technical issues. Cleaned sequencing reads of placental FcRs were then mapped to the human reference genome (assembly GRCh38, Gencode annotation release 25) using STAR to generate read counts for each of annotated genes. The raw gene read count data was normalized using edgeR (Robinson et al., 2010).

### FcR binding antibody multiplex assay

Fv and Fc profiles of plasma antibodies were characterized using a multiplexed Fc array assay as described previously (Brown et al., 2017; Brown et al., 2018). In brief, recombinant proteins were covalently coupled to fluorescent beads (Luminex). To display biotinylated peptides, streptavidin (Rockland) was coupled to beads first to capture the peptides. In duplicate, test samples were diluted 1:250 or 1:1,000 into a 384-well microplate (Greiner Bio One) containing ~500 beads of each specificity per well. Following bead opsonization, beads were washed and subsequently incubated with PE-conjugated Fc detection reagents, including human FcRs tetramers (Boesch et al., 2014), anti IgG (Southern Biotech), and C1q (Sigma-Aldrich), followed by a final wash. Data were acquired on a FlexMap3D instrument (Luminex) and raw data were reported as MFI values.

### Statistical analysis

The statistical analysis plan was finalized prior to data analysis. Multivariable linear regression models with sandwich variance estimates were used to evaluate the association between transfer efficiency and covariates of interest. Before regression, transfer efficiency was rank-gauss transformed (i.e. they are ranked and the ranks are transformed by the inverse Gaussian probability function). Missing antibody-specific transfer efficiencies were imputed using the R package mice. Principal component analysis of glycan markers was performed after the markers were scaled to 1. Global transplacental transfer was calculated for each patient and assigned a transfer score. An antigen-specific IgG transplacental transfer score was assigned to each distinct IgG specificity. All statistical procedures were implemented using the R language and environment for statistical computing and graphics. False discovery rate (FDR) p values are reported as FDR values.

## Acknowledgements

We thank Erin McGuire for technical support. We also thank the study participants and investigators from the WITS and CHAVI009 cohorts. The authors thank the children and families for their participation in Pediatric HIV/AIDS Cohort Study (PHACS), and the individuals and institutions involved in the conduct of PHACS. D.R.M., is supported by an American Society of Microbiology Robert D. Watkins Graduate Research Fellowship, a Burroughs Wellcome Graduate Diversity Fellowship, and an NIH NIAID Ruth L. Kirschstein National Research Service Award F31 F31AI127303. Y.F., is supported by NIH NIAID R01 AI122991. X.P., is supported by NIH NIAID R21AI125040 and R21AI120713. This work was partially funded through an R03 from NICHD to G.G.F., (HD085871). G.G.F., and S.R.P., are partially supported by IMPAACT. Overall support for the IMPAACT group is provided by the National Institute of Allergy and Infectious Diseases (NIAID) (U01 AI068632) and the Eunice Kennedy Shriver National Institute of Child Health and Human Development (NICHD). S.R.P., is supported by NIH NIAID DP2 HD075699, R01 AI106380, and P01 AI117915. The Pediatric HIV/AIDS Cohort Study (PHACS) was supported by the Eunice Kennedy Shriver National Institute Of Child Health & Human Development (NICHD) with co-funding from the National Institute Of Dental & Craniofacial Research (NIDCR), the National Institute Of Allergy And Infectious Diseases (NIAID), the National Institute Of Neurological Disorders And Stroke (NINDS), the National Institute On Deafness And Other Communication Disorders (NIDCD), Office of AIDS Research (OAR), the National Institute Of Mental Health (NIMH), the National Institute On Drug Abuse (NIDA), and the National Institute On Alcohol Abuse And Alcoholism (NIAAA), through cooperative agreements with the Harvard T.H. Chan School of Public Health (HD052102) and the Tulane University School of Medicine (HD052104).

Note: The conclusions and opinions expressed in this article are those of the authors and do not necessarily reflect those of the National Institutes of Health or U.S. Department of Health and Human Services.

## Author Contributions

D.R.M., G.G.F., and S.R.P. conceptualized and conceived the study. D.R.M., S.H.L., F.Y., J.A.W., E.A.H., R.G., J.F.M., and M.J. performed experiments and analyzed the data with help from Y.F., G.A., M.E.A., X.P., G.G.F., S.R.P.. G.S. provided critical samples for the study. All authors discussed the results and commented on the manuscript. Funding acquisition: D.R.M., X.P., G.G.F., and S.R.P.

---

**Table S1.**
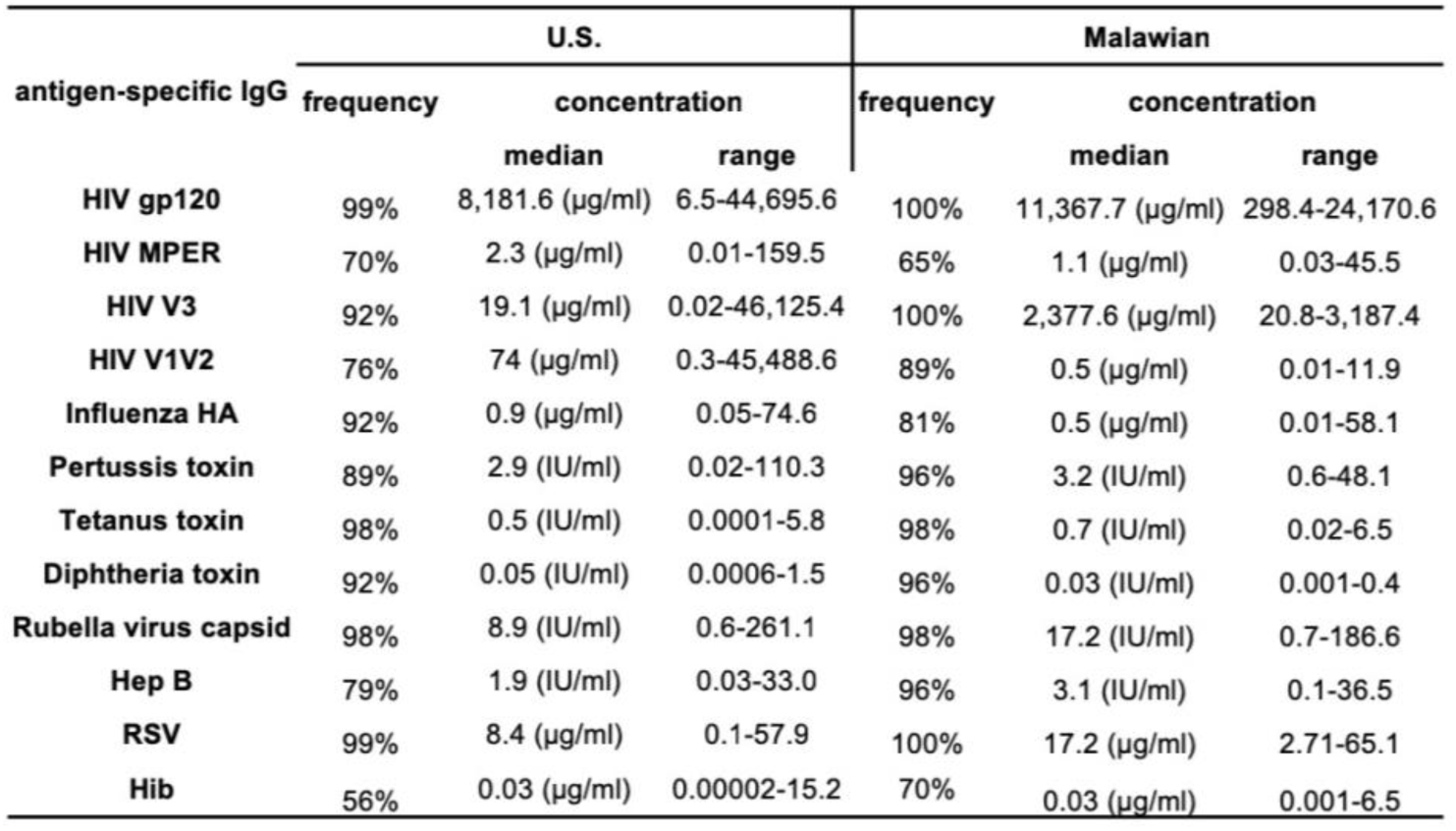
Frequency and concentrations of HIV and non-HIV pathogen-specific IgG response in U.S. and Malawian HIV- infected women.

**Table S2.** Antigen-specific IgG glycan Principal Components (PC) percent variance in U.S. HIV-infected women with variable transplacental IgG transfer.

| IgG Fc Glycan Components | gp120-specific IgG | tetanus toxoid-specific IgG | pertussis-toxin-specific IgG |
| --- | --- | --- | --- |
| **PC 1:** G0, G1, G2, mono-sialic acid, and total sialic acid | **64<sup>a</sup>** | **62** | **59** |
| **PC 2:** fucose, disialic acid, and bisected | **24** | **16** | **19** |
| PC 3 | 5 | 11 | 11 |
| PC 4 | 4 | 7 | 7 |
| PC 5 | 2 | 3 | 2 |
| PC 6 | 1 | 0 | 2 |
| PC 7 | 0 | 0 | 0 |
| PC 8 | 0 | 0 | 0 |

<sup>a</sup> % variance Bolded values indicate variance >20%

**Figure S1:**
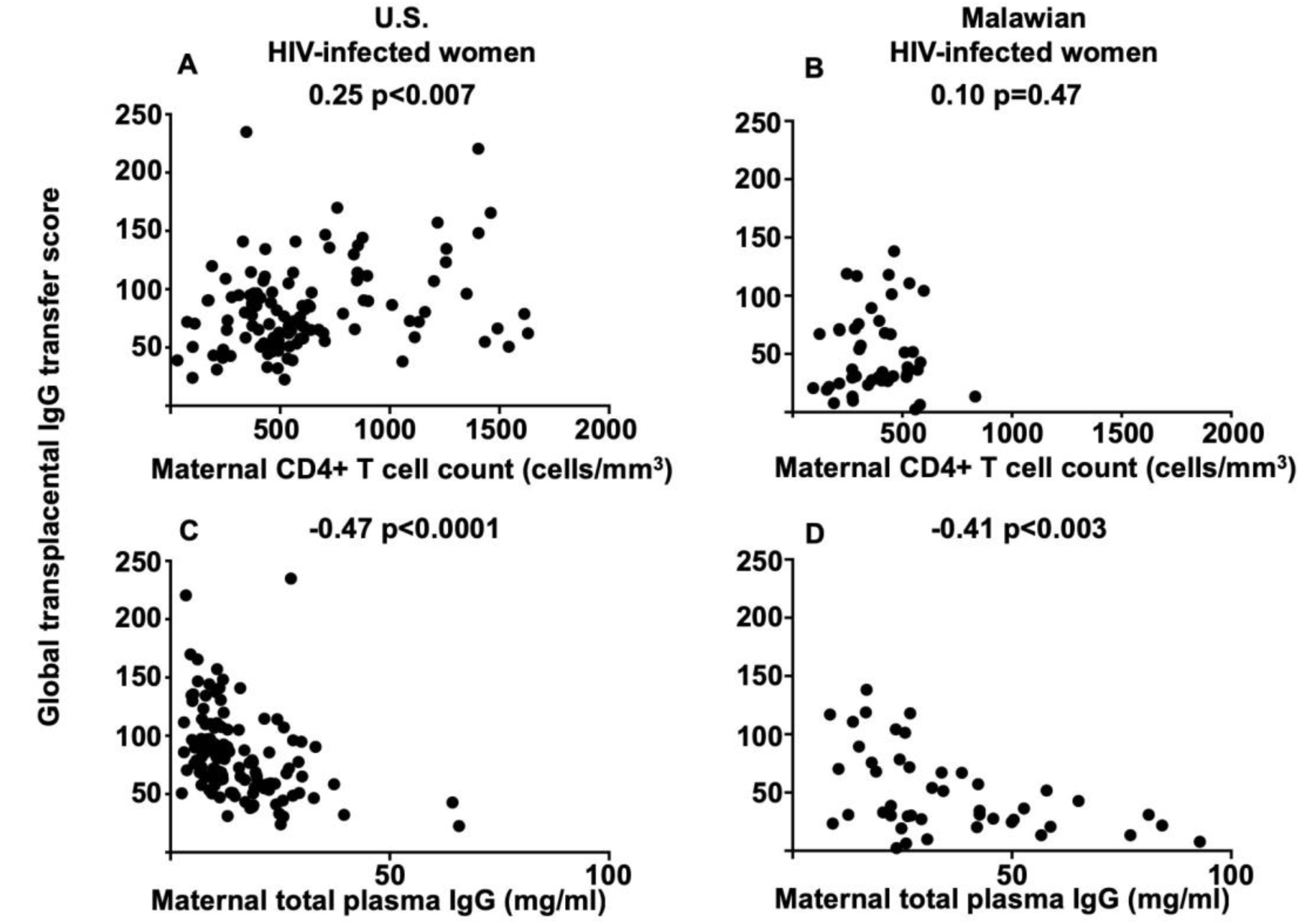
Maternal CD4+ T cell counts and total plasma IgG concentrations and transplacental IgG transfer efficiency. Correlation of maternal CD4+ T cell counts in A) U.S. and B) Malawian HIV-infected women and transplacental IgG transfer efficiency. Correlations of maternal total plasma IgG concentrations in C) U.S. and D) Malawian women and transplacental IgG transfer efficiency. Spearman correlation and p values are shown.

**Figure S2:**
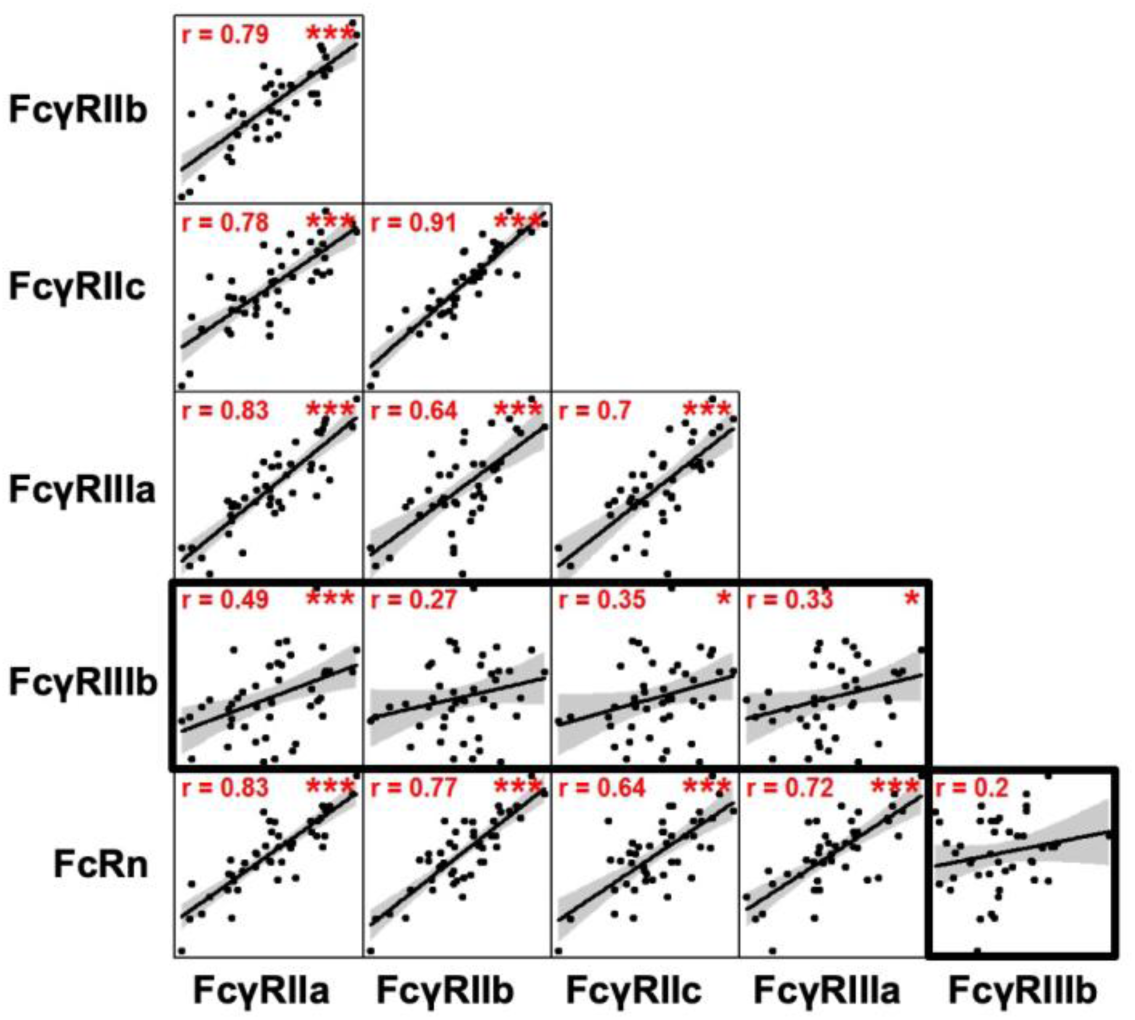
Strong correlations between the levels of mRNA expression of FcRs in placental tissues of HIV-infected Malawian women. Fc receptor expression levels in the placenta highly correlated between FcRn, FcyRMa, FcyRllb, FcyRNc, FcyRMIa (r: 0.64-0.91), yet there was a low correlation of FcyRlllb and other FcRs (r: 0.2-0.49, bolded boxes) in the placenta. Line of best fit is shown and the 95. pointwise confidence interval is shown in grey. * denotes medium correlation and *** denotes strong correlation.

**Figure S3:**
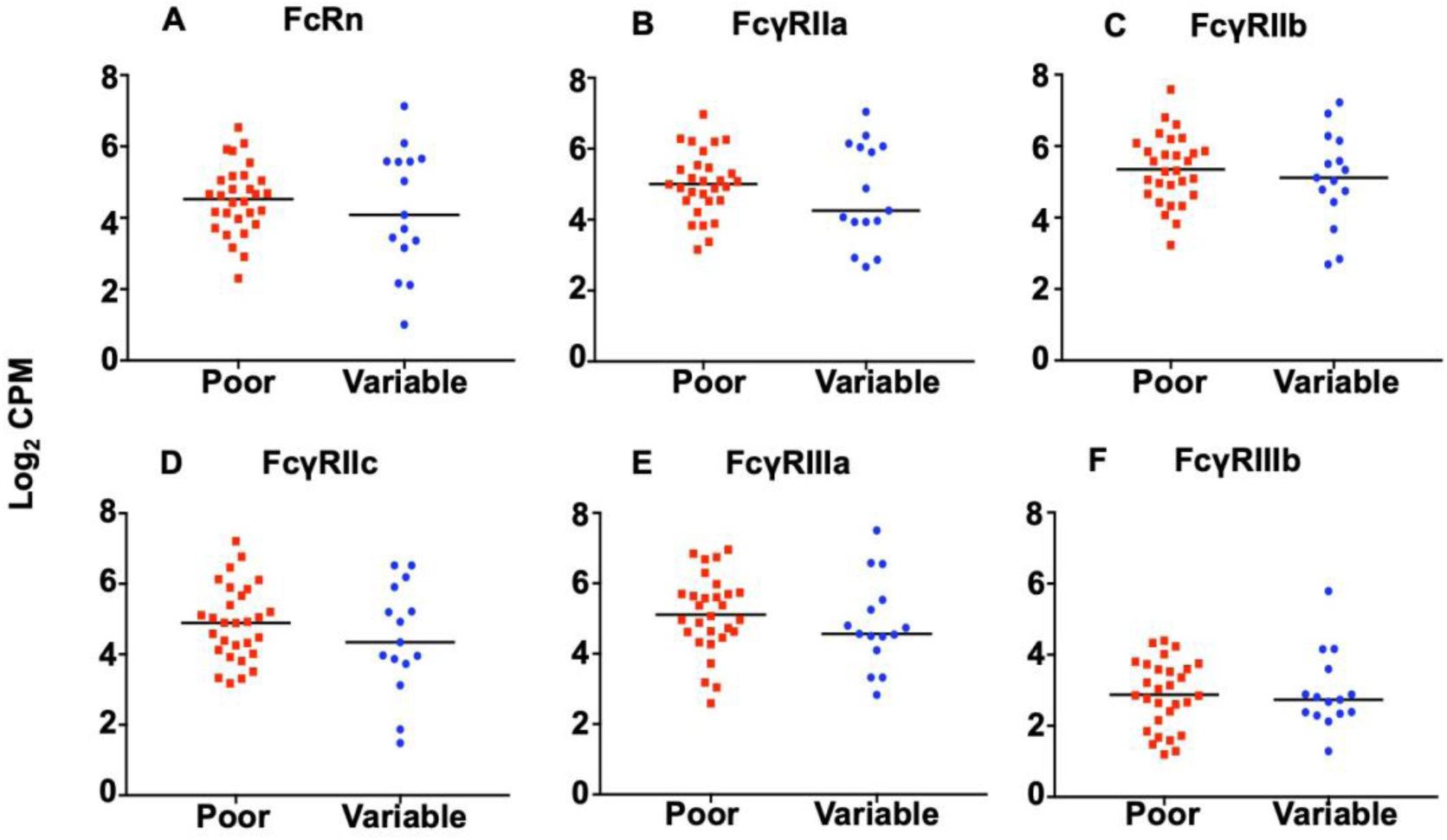
No difference in placental Fc receptor expression levels in HIV-infected Malawian women with distinct transplacental IgG transfer phenotypes. Placental mRNA expression levels of (A) FcRn, (B) FcyRlla, (C) FcyRllb, (D) FcyRllc, (E) FcyRllla, and (F) FcyRlllb in HIV-infected women with poor transplacental IgG transfer efficiency (red squares) and variable transplacental IgG transfer efficiency (bluecircles). Log_2_CPM denotes the counts per million of RNA copies. All comparisons of FcR expression between women with poor and variable placental transfer were not statistically significant (raw p>0.05).

**Figure S4:**
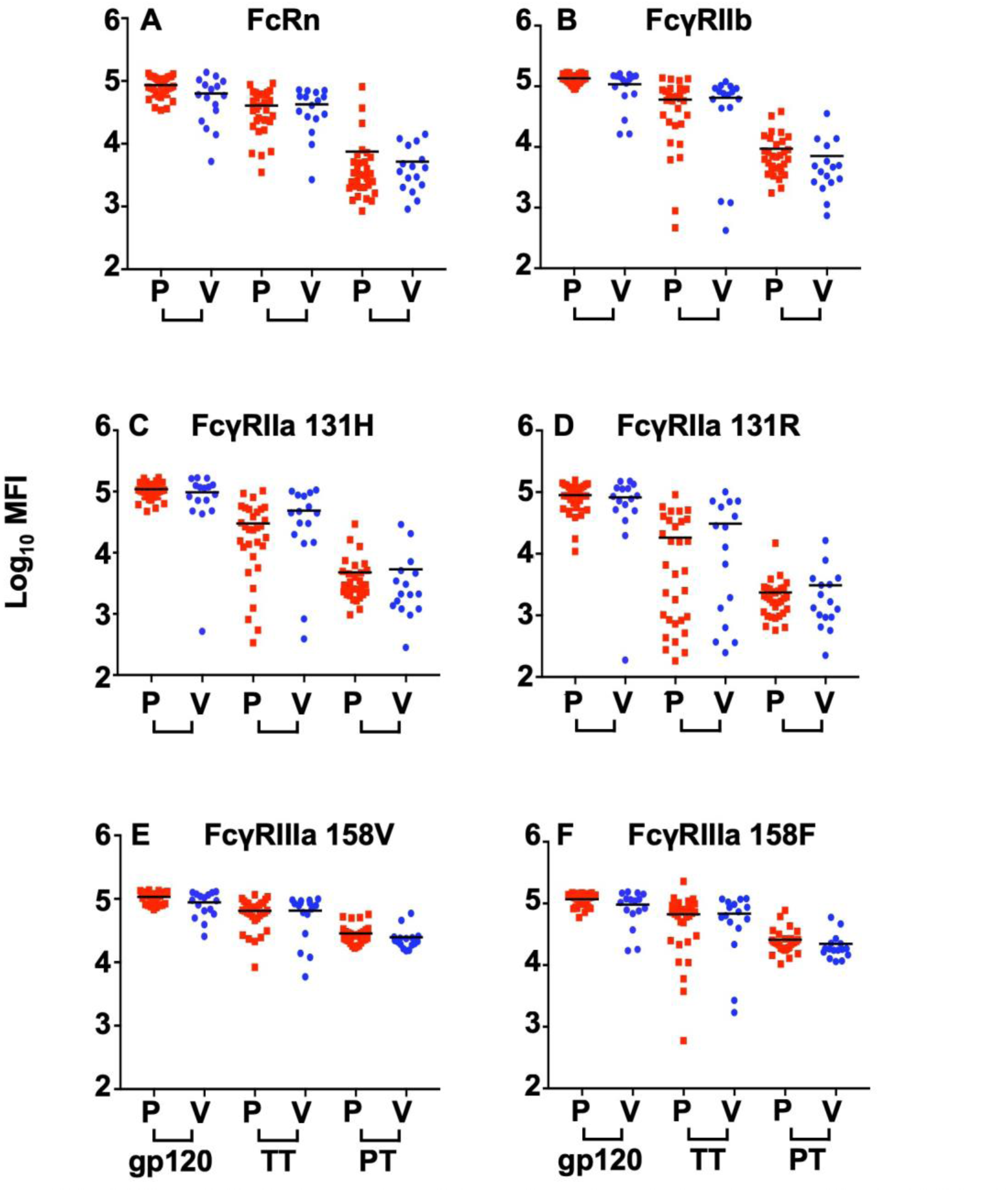
HIV gp120, tetanus toxoid, and pertussis toxin-specific IgG binding toplacental FcRs in HIV-infected Malawian women with poor (P) and variable (V)placental IgG transfer. Maternal gp120, tetanus toxoid (TT), and pertussis toxin (PT)- specific IgG binding to (A) FcRn, (B) FcyRllb, (C) FcyRlla 131H, (D) FcyRlla 131R, (E)FcYRIIIa 158V, and (F) FcyRllla 158F. FcyRlla 131H and FcYRIIIa 158V are high affinity IgG binding FcRs, whereas FcyRlla 131R and FcyRllla 158F are lower affinity IgG binding FcR polymorphisms. HIV-infected women with poor (P) transplacental IgGtransfer efficiency are shown in red squares and variable (V) transplacental IgG transfer efficiency in blue circles. Log10 MFI denotes the mean fluorescent intensity.

**Figure S5:**
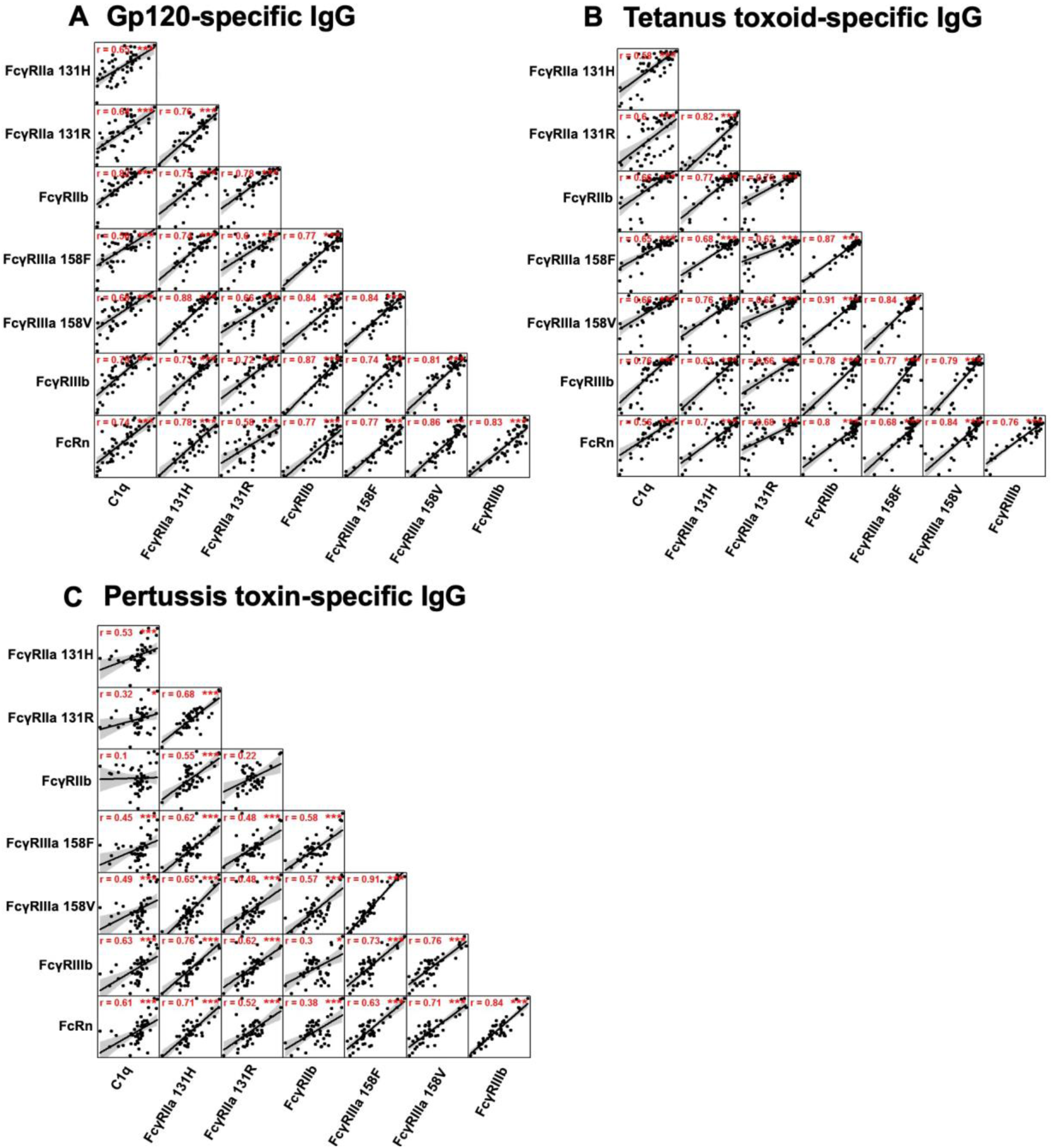
Correlations of binding magnitude of gp120, tetanus toxoid, andpertussis-specific IgG to distinct placental FcRs. Correlation of maternal gp120,tetanus toxoid, and pertussis toxin-specific IgG binding to C1q and placentallyexpressed FcRn, FcyRllb, FcyRlla 131H, FcyRlla 131R, FcyRllla 158V, FcyRllla158F, FcyRlllb in HIV-infected Malawianwomen. Line of best fit is shown and the 95. pointwise confidence interval is shown in grey.

**Figure S6:**
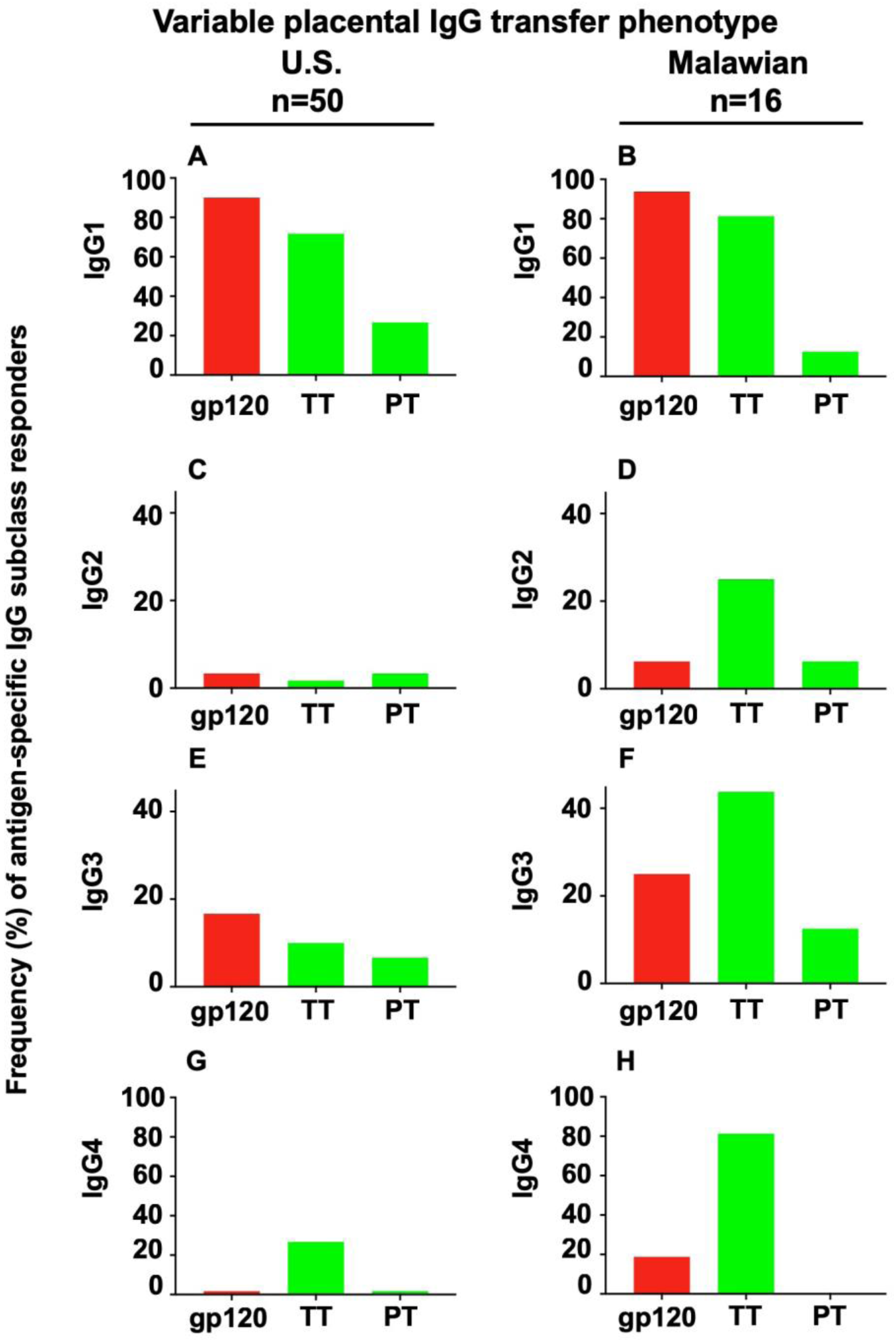
Frequency of gp120, tetanus toxoid (TT), and pertussis toxin(PT)-specific IgG subclass responders among HIV-infected, U.S. andMalawian women with variable transplacental IgG transfer. Frequency ofplasma gp120, tetanus toxoid (TT), and pertussis toxin (PT)-specific lgG1 (A,B), lgG2 (C, D), lgG3 (E, F), lgG4 (G, H) subclass responses in U.S. (A, C, E,G) and Malawian (B, D, F, H) women with variable transplacental IgG transfer.

